# CelLink: integrating single-cell multi-omics data with weak feature linkage and imbalanced cell populations

**DOI:** 10.1101/2024.11.08.622745

**Authors:** Xin Luo, Yuanhao Huang, Yicheng Tao, Fan Feng, Alexander Hopkirk, Thomas S.R. Bate, Diane C. Saunders, Peter Orchard, Catherine Robertson, Shristi Shrestha, Jean-Philippe Cartailler, Stephen C. J. Parker, Marcela Brissova, Jie Liu

## Abstract

Single-cell multi-omics technologies capture complementary molecular layers, enabling a comprehensive view of cellular states and functions. However, integrating these data types poses significant challenges when their features are weakly linked and cell population sizes are imbalanced. Currently, no method efficiently addresses these two issues simultaneously. Therefore, we developed CelLink, a novel single-cell multi-omics data integration method designed to overcome these challenges. CelLink normalizes and smooths feature profiles to align scales across datasets and integrates them through a multi-phase pipeline that iteratively employs the optimal transport algorithm. It dynamically refines cell-cell correspondences, identifying and excluding cells that cannot be reliably matched, thus avoiding performance degradation caused by erroneous imputations. This approach effectively adapts to weak feature linkage and imbalanced cell populations between datasets. Benchmarking CelLink on scRNA-seq and spatial proteomics datasets, as well as paired CITE-seq data, demonstrates its superior performance across various evaluation metrics, including data mixing, cell manifold structure preservation, and feature imputation accuracy. Compared to state-of-the-art methods, CelLink significantly outperforms others in imbalanced cell populations while consistently achieving better performance for balanced datasets. Moreover, CelLink uniquely enables cell subtype annotation, correction of mislabelled cells, and spatial transcriptomic analyses by imputing transcriptomic profiles for spatial proteomics data. CelLink sets a new milestone for multi-omics data integration. Its great ability to impute large-scale paired single-cell multi-omics profiles positions it as a pivotal tool for building single-cell multi-modal foundation models and advancing spatial cellular biology.

## Introduction

The integration of single-cell multi-omics data provides complementary information from various molecular layers for individual cells. This approach enhances our understanding of cellular functions, states, and interactions by capturing diverse biological signals such as epigenomic, transcriptomic, and proteomic data, each offering unique insights into cellular processes. Despite its promise, the integration of multi-omics data still faces *two challenges*. The first challenge is handling proteomics data with its rapid expansion, which exhibits weak feature overlap and distinct systematic biases compared to other data modalities. For example, the relationship between mRNA and protein expression levels may show a weak correlation due to multiple factors, including variations in their respective degradation rates and the influence of post-transcriptional regulatory mechanisms [1]. The second challenge is integrating multi-omics data with significant discrepancies in cell population sizes across different data modalities. Imbalanced cell populations between datasets often arise from the inherent characteristics of the datasets and the varying data quality control standards applied during preprocessing. For example, for a donor from the Human Pancreas Analysis Program [2] (i.e., HPAP024), the scRNA-seq data contains 54 beta cells comprising 4.26% of the total while the CODEX data includes 35,140 beta cells representing 14.49% of its total after strict data quality control.

Traditionally, methods such as Seurat V3 [3] and LIGER [4] require significant overlap in features between datasets, such as scRNA-seq with scATAC-seq or with spatial transcriptomics. These methods are not suitable when the datasets have *weak feature correspondence*. To address this, MMD-MA [5] aligns the cell-cell similarity (manifold) structures between the two data types after projecting them into the kernel spaces. SCOT v2 [6] aligns unpaired multi-omics data by comparing the distance between samples rather than comparing the samples themselves across modalities via Gromov-Wasserstein optimal transport. MaxFuse [7] progressively updates the cell-cell matching between the two data types through fuzzy smoothing. Nonetheless, it is common for single-cell multi-omics datasets from the same tissue to have dramatically different cell compositions, especially when different quality control standards are utilized to process the datasets. None of these methods perform well on datasets with imbalanced cell population sizes.

To overcome these challenges, we developed CelLink, a novel method to integrate single-cell multi-omics data. CelLink adeptly handles weak feature linkage through normalization and smoothing techniques, effectively neutralizing systematic biases introduced by different biotechnologies. Furthermore, it employs a dynamic cell-cell correspondence framework, which is iteratively updated to capture the imbalanced cell populations in datasets, resulting in satisfactory performance in case of discrepant cell populations. Meanwhile, CelLink demonstrates a slight superiority in scenarios of balanced populations. The benchmarking results show that CelLink consistently outperforms other state-of-the-art integration methods across a collection of datasets under various evaluation metrics. Moreover, CelLink is the only method that automatically identifies cells that cannot be aligned between the multi-omics datasets. This avoids the issue where including such cells would degrade the alignment performance, as the imputed feature values for these cells would inherently be problematic.

Additionally, CelLink has *three unique utilities* after integration. The first is cell subtyping and mislabeled cell identification. The precision for cell annotation is crucial for accurate treatment and prognosis. For example, distinguishing accurately between acute lymphoblastic leukemia (ALL) [8] and acute myeloid leukemia (AML) [9] subtypes is essential, as each subtype requires a different treatment regimen, directly affecting patient outcomes. The second is comprehensively analyzing spatial gene expression patterns and cell-cell communications. This capability is significant as current biotechnologies cannot simultaneously measure all gene expressions and spatial coordinates for single cells. For instance, technologies like CosMx [10] measures only a few thousand genes, significantly less than the 20,000 coding genes in the human genome, limiting their utility for complete spatial transcriptional profiling. The third is the precise feature imputation across multi-omics data. By simulating large-scale datasets with hundreds of thousands of samples and testing on CITE-seq datasets, CelLink robustly imputes missing features in datasets with varying feature correlations. Such large-scale feature imputation is pivotal for building single-cell multi-modal foundation models. These foundation models are in urgent need because of their significance in providing a comprehensive and interconnected understanding of disease mechanisms, like Type 2 Diabetes (T2D), across multiple biological layers from genomics to metabolomics.

## Results

### Cell-cell alignment between modalities via iterative optimal transport

The inputs of CelLink are the expression matrices of the two modalities *X* and *Y* (Fig. 1A). First, to reduce the discrepancies of the two data modalities, we normalize them on a common numeric scale. Next, to mitigate the inherent biases arising from different biotechnological platforms, we perform cell-wise smoothing from *k*-NN graphs based on the feature profile for each modality. Then, we define “linked features”, which refer to corresponding entities in *X* and *Y*, such as the GCG coding gene in scRNA-seq and the GCG protein marker in CODEX data. We denote the profiles of these linked features in *X* and *Y* as *X*_*l*_ and *Y*_*l*_, respectively. After preprocessing the datasets, we input the normalized and smoothed linked features 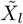 and 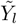 into the integration pipeline of CelLink. The integration pipeline offers two choices. If cell type or cell state is provided, it starts with Balanced Optimal Transport (BOT), continues with iterative unbalanced Optimal Transport (UOT) until convergence, and finally transfers the feature profile from one modality to another. Otherwise, it just performs a one-time interoperable UOT, followed by feature transferring.

**Figure 1:**
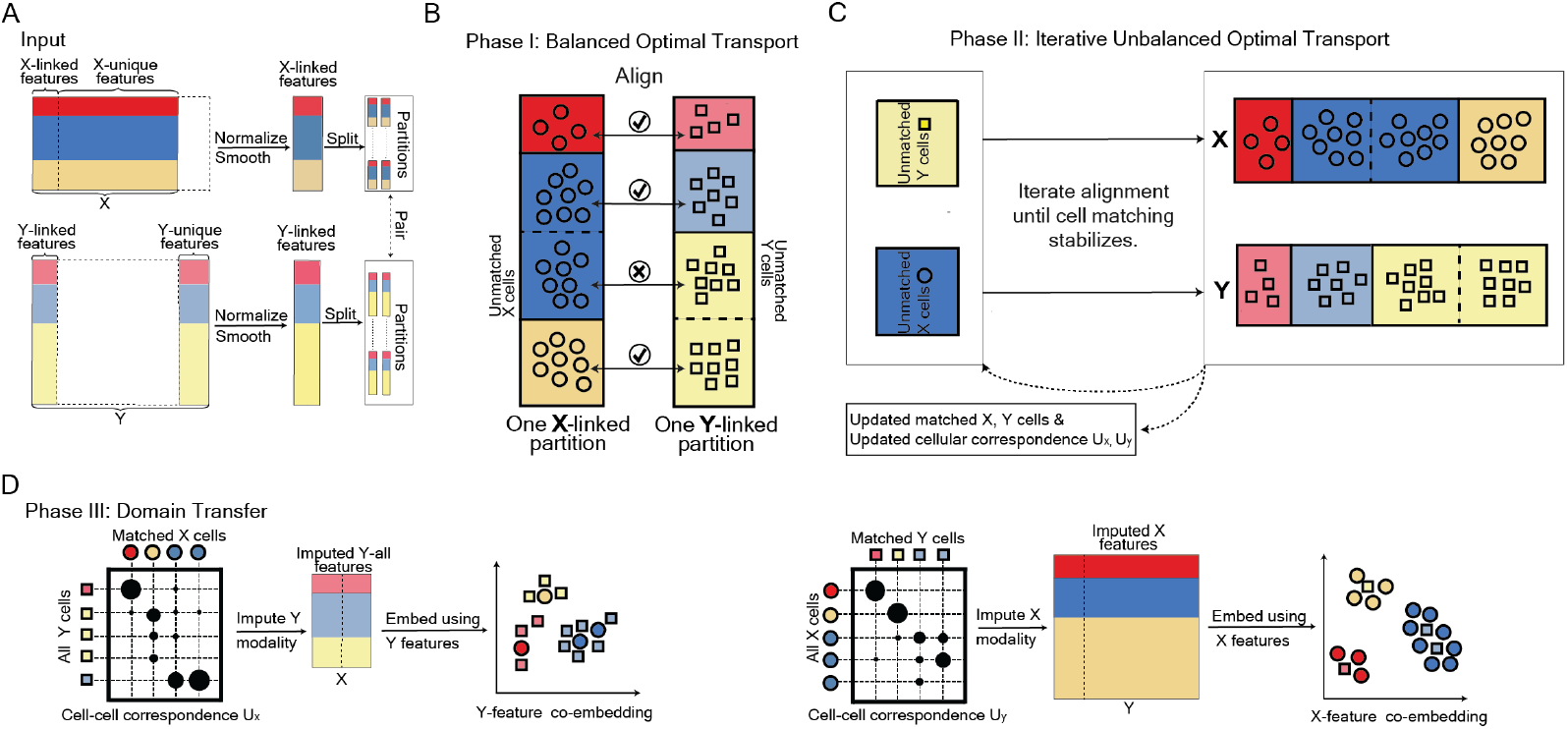
The pipeline of CelLink. **A.** The input is composed of the all-feature and linked-feature of X and Y modalities. CelLink utilizes the all-feature modality to smooth and denoise the linked-feature modality by the nearest neighbor (NN) graph and low-rank approximation. The smoothed linked modality *X*_*ss*_ and *Y*_*ss*_ are used to perform the integration. **B**. The first phase is to map cells using balanced optimal transport. CelLink predicts the cell type for each cell by the weight from the cell-cell transport map. The matched cells will be filtered while the unmatched ones will be re-aligned in the next phase. The transport map will be stored as the cell-cell correspondence matrix. **C**. The second phase is to iteratively align the unmatched cells using unbalanced optimal transport. The iterative alignment is performed separately for each modality. In each iteration, unmatched cells are identified via the cell-cell transport map and re-aligned in the next run. This transport map is then preserved as the corrected cell-cell correspondence matrix. The alignment stops when the predicted cell type for all unmatched cells does not change, indicating that they cannot be aligned and the model reaches convergence. **D**. In the third phase, all cell-cell correspondence matrices are consolidated to establish the final cell-cell correspondence framework. CelLink mutually imputes the feature profiles between modalities. The goodness of the alignment is evaluated after jointly embedding the imputed modalities. The output of CelLink is the final correspondence matrix *U*_*x*_, *U*_*y*_ and the imputed modalities.

In the first phase, BOT is performed to measure the cellular correspondence between modalities (Fig. 1B). To align cells fairly, we determine the mass of all cells in OT as 1.0. The cost matrix of BOT, defined as *D*_*xy*_, is constructed based on the pairwise dissimilarity between cells in *X* and *Y*. See the Methods section for more details. Given that BOT requires the total mass of cells equal across modalities, we split each data into multiple partitions, ensuring that the number of cells in each pair of partitions is close. The outcome of BOT is a transport map, which establishes the initial cell-cell correspondence between modalities in the iterative procedure, denoted as *U*_0_. When the cell-type or cell-state populations of the two data are significantly imbalanced, some cells will be mapped to erroneous cell types or cell states. This misalignment occurs because the total mass attributed to a particular cell type or cell state does not match between the two datasets. To solve this problem, we filter out all matched cells and retain all unmatched cells whose predicted cell types are erroneous, followed by correcting their alignments in the following phase. The predicted cell type is the one with the highest weight from the other modality calculated via *U*_0_.

In the second phase, we conduct iterative UOT for the unmatched cells in one modality to all cells in another (Fig. 1C). UOT does not require the total mass of cells to be equal between the two datasets, which largely addresses the challenge of disparate cell population sizes. However, the optimization of the optimal transport algorithm faces a trade-off between efficiency and global convergence issues. This results in incorrect matches for certain cells, whose feature distributions overlap with other similar cell types due to the weak feature linkage between datasets. To resolve this issue, we iteratively remove the correctly matched cells in each round and realign the remaining unmatched cells. This strategy gradually reduces the matching space for the remaining cells thereby improving their matching. The iteration continues until the predicted cell types for unmatched cells stabilize, indicating the convergence of UOT. This stability suggests that the remaining cells cannot be aligned with cells in the other modality. The cellular correspondence matrices in each iteration are assumed as *U*_1_, *U*_2_, …, *U*_*t*_, where *t* denotes the iteration times. The remaining unmatched cells will be excluded in the subsequent step.

In the third phase, we reciprocally transfer the feature profile from one modality to another. The outputs are the imputed feature profiles of all modalities along with their cell-cell correspondence matrices (Fig. 1D). The series of cellular correspondence matrices *U*_0_, *U*_1_, …, *U*_*t*_ is consolidated into a unified matrix *U* based on cell IDs, wherein the cell correspondence vector for each initially unmatched cell in *U*_0_ is substituted with its counterpart in *U*_*f*_. Here, *f* represents the index of the correspondence matrices that correct the unmatched cell. The feature transfer for one data is achieved by multiplying its consolidated correspondence matrix *U* with another data. After feature transfer, *X* and *Y* are jointly embedded into a low-dimensional space for further evaluation.

### Benchmarking CelLink in the integration of scRNA-seq and CODEX data

We benchmarked CelLink across three scRNA-seq and CODEX datasets. One of these datasets features a balanced cell population of 10,000 randomly selected cells from tonsil tissue [11], with 46 protein markers from CODEX data. The other two datasets from HPAP (Home Pancreas Analysis Program) donors exhibit imbalanced cell populations from pancreas tissue [2]. The first HPAP dataset includes 1,340 cells from scRNA-seq and 12,030 cells from CODEX data with 24 protein markers, while the second one includes 1,743 cells from scRNA-seq and 1,516 cells from CODEX data with 24 protein markers as well. The cell-type composition of these three datasets is shown in Supplementary Table 1-3. We benchmarked CelLink with four baseline methods: CelLink (non-iterative), MaxFuse [7], Seurat v3 [3], and SCOT v2 [6]. The non-iterative version of CelLink represents the case when no cell type or cell state is provided, prompting CelLink to perform the one-time interoperable UOT [12] in the second phase.

A key objective of multi-omics integration is to precisely align the feature profiles from one modality to another. Given the absence of paired cells in the scRNA-seq and CODEX data, we assessed the integration performance based on the quality of cell-type clustering and data-type mixing from five views.

Firstly, the UMAP plots, generated by embedding the original and imputed data into the shared latent space, intuitively showed that CelLink consistently mixed data modalities and separated cell types well with all kinds of feature profiles (Fig. 2A, Supp Fig. 1-3). Another method MaxFuse, although successfully integrated the dataset with balanced cell populations regarding the protein profile, failed to separate the cell types in datasets with imbalanced cell populations (Fig. 2A, Supp Fig. 1B, Supp Fig. 2B). Next, the preservation of local cohesion of data modalities and cell types was evaluated by the Graph Connectivity Score (GCS). This result indicated that CelLink perfectly conserved the local connection within cell types and data types using all kinds of feature profiles compared with other methods in all datasets (Fig. 2B, Supp Fig. 4A-4B). Subsequently, we assessed the preservation of global differences between data types and cell types using the Average Silhouette Width (ASW). CelLink significantly outperformed other methods in all datasets (Fig. 2C). Additionally, cell-type matching accuracy (CTMA) and the count of precisely matched cells (CPMC) are also considered critical metrics to determine whether a method can succeed in aligning the maximum number of cells. From the result, CelLink far surpassed other methods in the imbalanced datasets (Fig. 2D). The heatmaps revealed that CelLink is the only method capable of aligning different cell types with high purity in the presence of imbalanced cell population (Fig. 2E, Supp. Fig. 5A). Although CelLink naturally exhibits higher cell-type matching accuracy owing to its use of cell-type labeling, it does not promote cells of the same type to align. Instead, it just allows unmatched cells to realign until their matched cell types no longer change.

**Figure 2:**
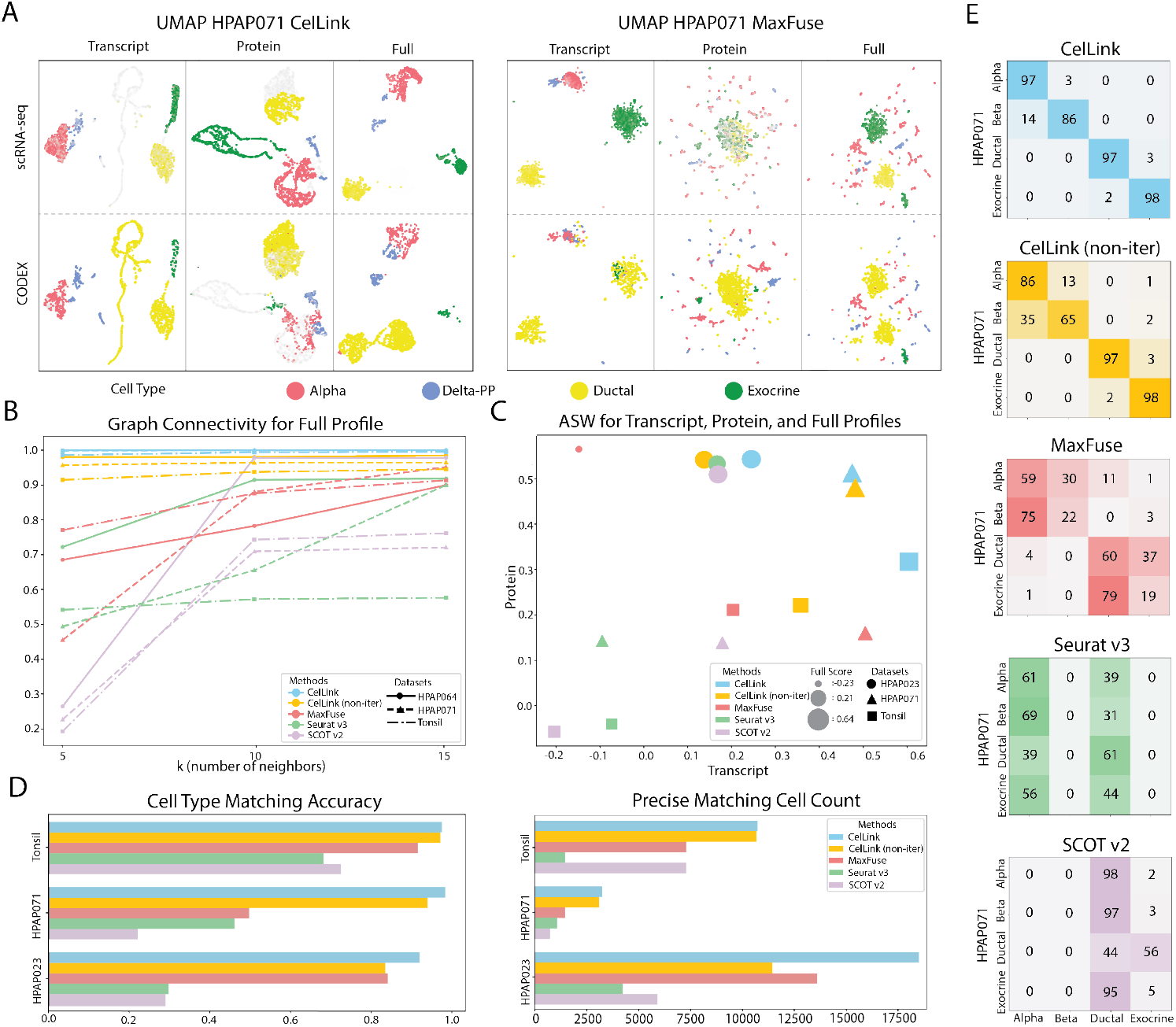
Systematic benchmarking of CelLink on scRNA-seq and CODEX datasets. **A.** The UMAP plots exhibit the intuitive integration results from CelLink and MaxFuse on a T1D HPAP071 donor with the transcript, protein, and full (transcript & protein) profiles. For the transcript profile, scRNA-seq data refers to the original profile while CODEX refers to the the imputed one. For the protein profile, scRNA-seq data refers to the imputed profile while CODEX refers to the original one. **B**. The line plot displays the graph connectivity scores for CelLink and four other baselines across three scRNA-seq and CODEX datasets. Each line indicates the trend of the score when the edges of the graph are built with the increasing number of neighboring cells. The color of the line represents the method. The line type and point shape represent the dataset. **C**. The scatterplot displays the average silhouette width for CelLink and four other baselines across three datasets using the transcript, protein, and full profiles. The size of the point corresponds to the ASW of the full profile. **D**. Left: The horizontal barplot exhibits the cell-type matching accuracy for CelLink and four other baselines across three datasets. Right: The horizontal barplot exhibits the number of cells that are precisely matched across three datasets. **E**. The heatmaps display the detailed cell-type matching result for CelLink and four other baselines on the HPAP071 dataset.

### Additional benchmarking CelLink in the integration of CITE-seq data

Since scRNA-seq and CODEX data do not contain paired cells, We additionally benchmarked CelLink on four CITE-seq datasets, which provide ground-truth transcriptomic and proteomic profiles for each cell. The first two datasets, derived from PBMC and BMC data [13], were sub-sampled to 10,000 cells each. The third dataset was generated by splitting the PBMC CITE-seq data into imbalanced groups: one group with 3,200 cells and another with 6,800 cells. Similarly, the fourth dataset was created by splitting the BMC CITE-seq data into imbalanced groups: one group with 8,000 cells and another with 2,000 cells. Detailed cell-type distributions of the third and fourth datasets are provided in Supplementary Table 4-5. For these two datasets, we used the transcriptomic profile from the first group and the proteomic profile from the second group for integration, while their corresponding alternative feature profiles served as the masked ground truth for evaluating the integration performance.

From the UMAP plots, CelLink achieved excellent mixing of data modalities and separation of cell types using the transcript, protein, and full feature profiles in all of the datasets (Fig. 3A-B, Supp Fig. 6A-B). In terms of the count of precisely matched cells and cell-type matching accuracy, CelLink consistently outperformed all other baseline methods in all datasets (Fig. 3C).

**Figure 3:**
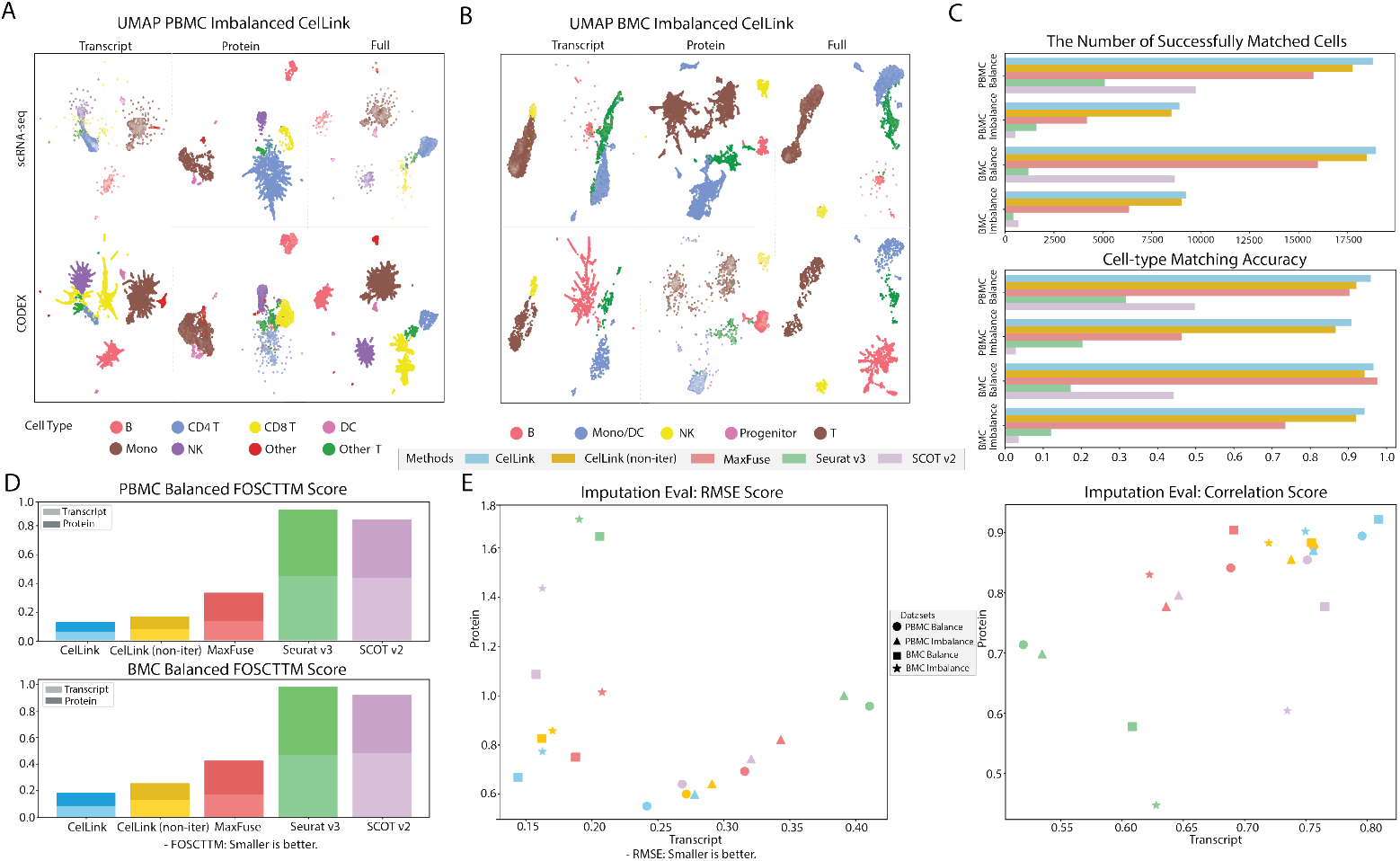
Systematic benchmarking of CelLink on CITE-seq datasets. **A.** The UMAP plots intuitively exhibit the integration results of CelLink on the imbalanced PBMC data with the transcript, protein, and full (transcript & protein) profiles. **B**.The UMAP plots intuitively exhibit the integration results of CelLink on the imbalanced BMC data with the transcript, protein, and full (transcript & protein) profiles. **C**. Top: The horizontal barplot displays the number of successfully matched cells for the five methods on four CITE-seq datasets. Bottom: The horizontal barplot displays the cell-type matching accuracy for the five methods on four CITE-seq datasets. **D**. Upper: The stacked barplot exhibits the FOSCTTM for the five methods on the PBMC CITE-seq dataset using the transcript and protein profile. Lower: The stacked barplot exhibits the FOSCTTM for the five methods on the BMC CITE-seq dataset using the transcript and protein profile. **E**. Left: The scatterplot manifests the RMSE score of the transcript and protein profiles for the five methods on all CITE-seq datasets. Right: The scatterplot manifests the Correlation score of the transcript and protein profiles for the five methods on all CITE-seq datasets.

With the ground-truth transcript and protein profiles for each cell, we induced three additional metrics to evaluate the integration performance at the single-cell level. The first metric, Fraction Of Samples Closer Than True Match (FOSCTTM), assesses how closely the true matching cell is aligned relative to other cells post-integration. CelLink achieved the best performance with both transcript and protein profiles in the PBMC and BMC CITE-seq datasets (Fig. 3D). The second metric, RMSE of Feature Imputation, calculates the average RMSE between the aligned and original feature profiles for all cells. CelLink consistently showed the lowest RMSE score across all datasets using both transcript and protein profiles (Fig. 3E). The third metric, Correlation of Feature Imputation, computes the average Pearson Correlation Coefficient between the aligned and original feature profiles. CelLink achieved the highest Correlation across all datasets using both transcript and protein profiles (Fig. 3E).

In conclusion, CelLink effectively recovers masked feature profiles after integration, demonstrating high precision in imputing both transcriptomic and proteomic features.

### CelLink enables cell subtyping and identifies mislabeled cells

While numerous cell-type annotation methods have been developed, efficient approaches for cell subtype annotation and correction are still lacking. CelLink not only enables annotating the subtypes of cells but also identifies the mislabelled cells.

We applied CelLink on this task using scRNA-seq and CODEX data from Bandyopadhyay, Shovik, et al. [14] of bone marrow tissue (Fig. 4A). We randomly subsampled 10,000 cells from scRNA-seq and CODEX data according to the provided cell lineage and subtype annotations (Fig. 4B, Supplementary table 6-7). With cell lineage, CelLink successfully integrated the scRNA-seq and CODEX data, as cells originating from the same lineage are grouped in the embedding space (Fig. 4C).

**Figure 4:**
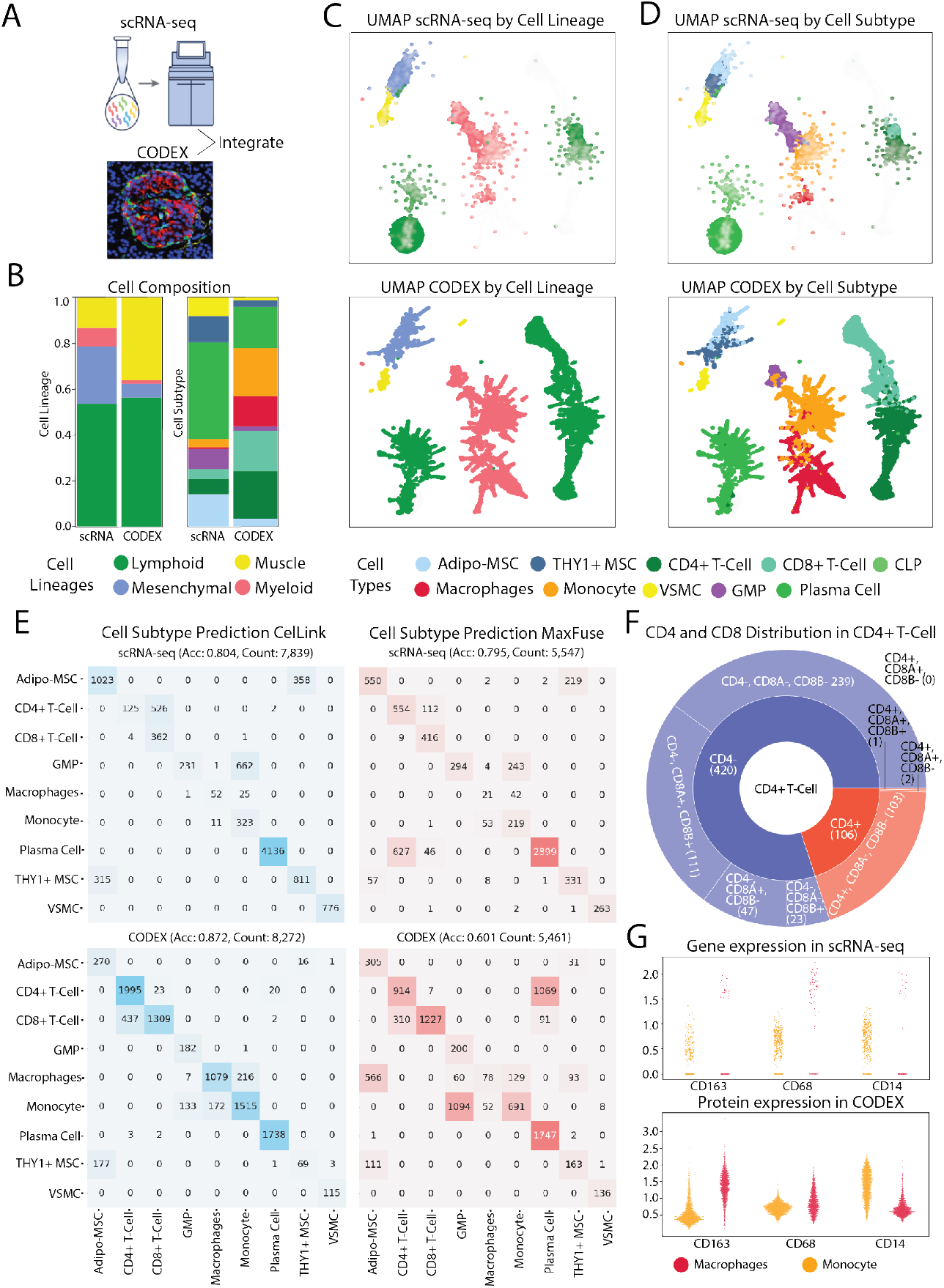
CelLink enables annotation and identification of cell subtypes. **A.** Overview of integrating scRNA-seq and CODEX data. **B**. The stacked plots depict the imbalanced cell composition between scRNA-seq and CODEX data grouped by cell lineage and cell subtypes, respectively. **C**. The UMAP plots depict the distribution of cells annotated by the cell lineage after integration in the embedding space. The four colored cell lineages are “Lymphoid”, “Muscle”, “Mesenchymal”, and “Myeloid”. **D**. The UMAP plots depict the distribution of cells annotated by the cell subtype after integration in the embedding space. The subtypes of Lymphoid cells are “CD4+ T-Cell”, “CD8+ T-Cell”, “CLP (Common Lymphoid Progenitor)”, and “Plasma Cell”. The subtypes of Mesenchymal cells are “Adipo-MSC” and “THY1+ MSC”. The subtypes of Myeloid cells are “Macrophages” and “Monocyte”. The subtype of the muscle cell is the “VSMC (Vascular Smooth Muscle Cells)”. **E**. The heatmaps exhibit the prediction performance of cell subtypes by CelLink compared with MaxFuse. Acc: The cell-subtype prediction accuracy. Count: The total number of cells whose subtypes are predicted precisely across all subtypes. **F**. The sunburst chart displays the distribution of CD4, CD8A, and CD8B coding genes among the mispredicted CD4+ T Cells. Each partition describes the number of cells with a distinct combination of CD4 and CD8 marker expressions. **G**. The strip plots display the distribution of CD163, CD68, and CD14 markers within Macrophage and Monocyte cells. The upper plot represents the scRNA-seq data, while the lower plot corresponds to the CODEX data.

Meanwhile, different cell subtypes were separated well in the result (Fig. 4D). Then, we predicted the cell subtypes in the CODEX data using the cell subtype annotation from the scRNA-seq data, and vice versa. Specifically, the subtype of each cell was predicted by the subtype with the highest weight in the inferred cell-cell correspondence matrix. The performance of subtype prediction was benchmarked with multiple baseline methods, including CelLink (non-iterative), MaxFuse, Seurat v3, and SCOT v2 [7, 3, 6]. CelLink significantly outperformed all other methods regarding the subtype prediction accuracy rate and the count of precisely matched cells (Fig. 4E, Supp Fig. 7A-B). Notably, CelLink achieved a high subtyping accuracy of 87.2% for the CODEX data, confirming its effectiveness in cell subtyping. However, a few subtypes were still not predicted perfectly, including CD4+ T-Cell, GMP, Adipo-MSC, and Macrophages in scRNA-seq and CD8+ T-Cell, Thy1+ MSC in CODEX. To investigate the reason, we examined the expression of marker genes in the mispredicted cells. Among the 526 mispredicted CD4+ T cells, only 106 expressed the CD4 gene while lacking expression of both CD8A and CD8B genes (Fig. 4F). This suggests that about 80% of the CD4+ T cells were incorrectly predicted as CD8+ T cells due to problematic annotation. Similarly, 267 out of 358 mispredicted Adipo-MSCs express Thy1, while 187 out of 315 mispredicted Thy1+ MSCs did not express Thy1, leading to cross-misprediction between Adipo-MSCs and Thy1+ MSCs (Supp. Fig. 7C). GMP cells were also primarily mispredicted for the same reason, with only 120 out of 662 mispredicted GMP cells expressed both CD34 and CD38 (Supp. Fig. 7C). Some Macrophage cells were mispredicted not only because of the wrong annotation but also due to their overlapping distribution with Monocyte cells regarding both gene and protein expressions (Fig. 4G). Although CD14 is the marker of Monocyte cells, and CD68 and CD163 are markers of Macrophage cells, they can still express each other’s markers at lower levels.

In summary, the misprediction of these subtypes is mainly due to two reasons, namely incorrect cell-subtype annotation in scRNA-seq, and the non-distinguishable expression level of the gene expression and protein marker intensities between specific subtypes. CelLink is the first method that enables accurate prediction and effectively identifying mislabeled cells.

### CelLink reveals spatial gene expression patterns and comprehensive cellular communications

Recently, there has been a fast-growing number of spatial proteomics datasets, such as CODEX, MIBI-TOF, IMC, and CosMx SMI [15, 16, 17, 18], which measure 30-100 protein markers for single cells along with their spatial coordinates. However, these proteomic datasets are limited in advanced spatial analyses like cell-cell communications due to the small number of measured markers. Additionally, current spatial transcriptomics technologies fall short of providing complete spatial transcriptomic profiling for single cells. For instance, CosMx [10] can measure only a few thousand genes, and 10x Visium [19] is limited to capturing gene expression data from predefined spots rather than single cells. CelLink addresses this limitation by imputing gene expression profiles for spatial proteomics data.

We applied CelLink to infer the spatial cell-cell communication in human pancreas tissue and validated the results by GLKB. We utilized a scRNA-seq and CODEX data from an AAB (auto-antibody positive) HPAP024 donor [2], with the scRNA-seq data comprising 1,266 cells, and the CODEX data comprising 12,351 cells (Supplementary table 8). To infer spatial cellular communication, we experimented with three steps. Firstly, CelLink integrated these two datasets and imputed 10,000 HVGs for the CODEX data. (Fig. 5A). To validate the imputation, we compared the spatial patterns of cell-type specific protein markers with their corresponding coding genes and found them all consistent (Supp Fig. 8A-D). Cells in the CODEX data are annotated with four cell types, and each cell type exhibited unique spatial patterns (Fig. 5B). Secondly, We used CellChat v2 [20] to identify significant ligand-receptor interactions between cell-type pairs, based on their expression levels and spatial proximity (see Methods). We summarized the results for several key ligand-receptor pairs in the pancreatic tissue (Fig. 5C). For example, beta cells act as the sender for the INS-INSR ligand-receptor pair and the IGF signaling pathway. Alpha and beta cells function as the sender and receiver for the GCG-GCGR, and GCG-GIPR ligand-receptor pairs. Thirdly, to validate these findings, we employed RAG (Retrieved Augmented Generation) of GLKB (Genomic Literature Knowledge Base) to search for supporting evidence from existing literature [21]. The retrieved answers from GLKB proved that all of the inferred cell-type level ligand-receptor interactions are correct (Fig. 5D).

**Figure 5:**
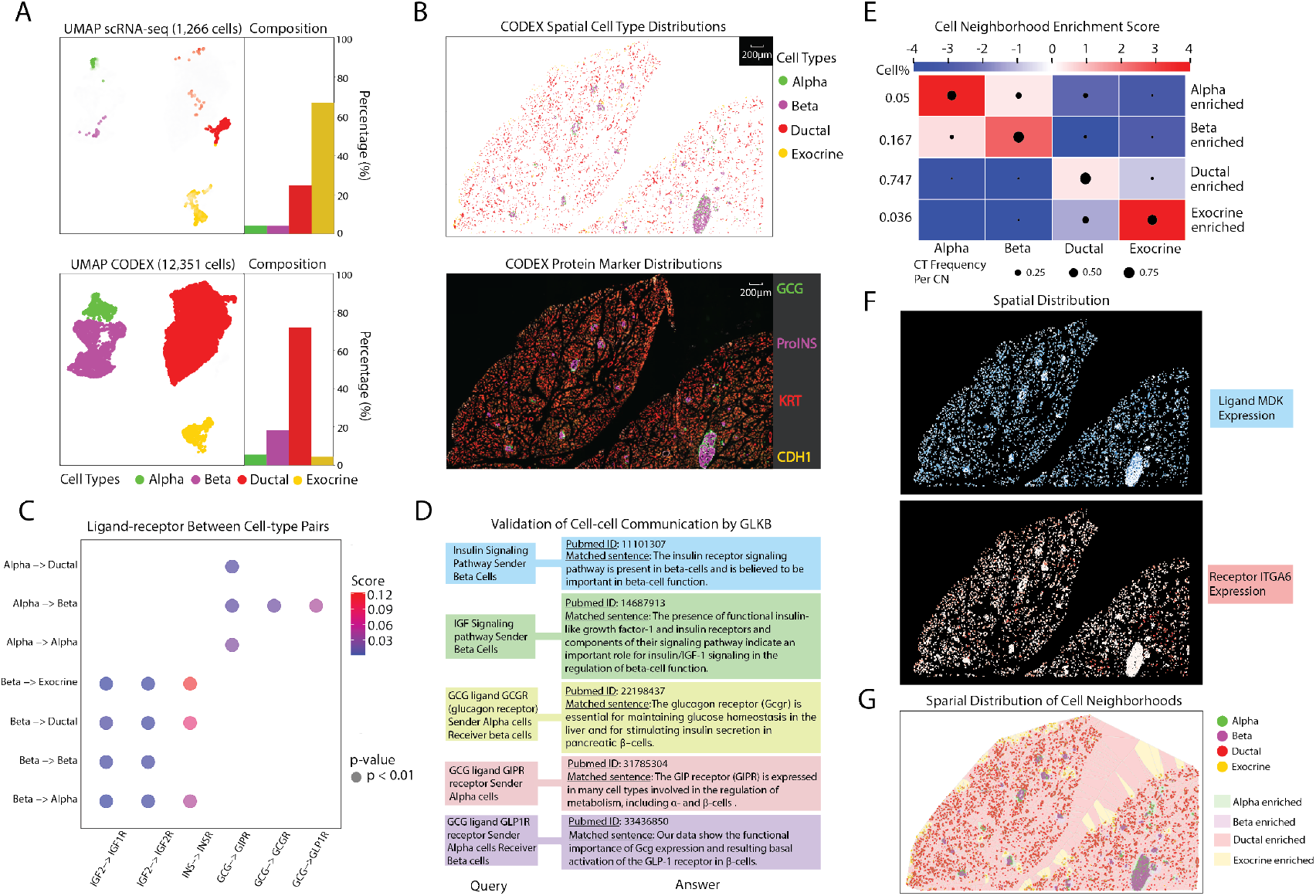
CelLink infers spatial cellular interactions in human pancreas tissue. **A.** Left: The UMAP plots depict the cell distribution from scRNA-seq and CODEX data colored by cell type after integration in the embedding space. Right: The barplots depict the imbalanced cell-type compositions between the 1,266 cells from scRNA-seq data and 12,351 cells from CODEX data. **B**. Upper: The spatial distribution of the cell types from CODEX data after cell segmentation within the sub-region of the head of the human pancreas tissue. Lower: The spatial distribution of the cell-type specific protein markers from the CODEX image within the sub-region of the head of the human pancreas tissue. **C**. The bubble plot demonstrates several significant and pivotal ligand-receptor interactions between cell-type pairs in the pancreas tissue. **D**. The figure illustrates the validation of the inferred cell-type resolution ligand-receptor interactions and signaling pathways in Figure C by RAG of GLKB. The left side displays the query content and the right side shows the retrieved answers from academic literatures in PubMed. **E**. The heatmap depicts the identified cell neighborhoods among the selected region in the CODEX data. The size of the point in the heatmap represents the percentage of a cell type within a specific cell neighborhood type. **F**. The spatial distribution of Ligand-receptor pair MDK-ITGA6 in the tissue. **G**. The spatial distribution of cell neighborhoods in the tissue.

To investigate the activities in different regions and microenvironments, we conducted cell neighborhood analysis for the CODEX data, identifying four distinct cell neighborhoods using CNTools [22]. Each cell neighborhood is dominated by one primary cell type mixed with others (Fig. 5E). The result indicates that alpha and beta cells are closely located in the spatial region, corresponding to human islets. In contrast, a subset of ductal cells is positioned near exocrine cells, consistent with the exocrine region of the tissue. Further, we detected the spatial distribution of ligand-receptor pairs regarding cell neighborhoods. For example, in the MDK-ITGA6 ligand-receptor pair, the sender cells are located in alpha and ductalenriched neighborhoods, while the receptor cells are found in ductal and exocrine-enriched neighborhoods (Fig. 5F-G). This result suggests a coordinated function among various cell neighborhoods, promoting the migration and adhesion of immune cells across the islet in response to the tissue damage and inflammation [23].

### Systematic Evaluating CelLink on its applicability through large-scale simulations

CelLink robustly integrates single-cell multi-omics datasets, whether the linkage between distinct modalities is weak or strong, and it demonstrates excellent scalability for large-scale datasets. To validate its adaptability and efficiency, we systematically evaluated CelLink using large-scale simulated data generated by a custom-built simulator. CelLink leverages the optimal transport algorithm, which does not rely on prior assumptions about input data distributions. Consequently, we generated datasets from normal distributions without loss of generality. See the Methods section for more details.

We tested CelLink on datasets with 100 linked-features whose correlations range from 0 to 0.9 (Supp. Fig. 9A). CelLink successfully integrated datasets with 10,000 samples across all levels of feature correlations except at correlation = 0. This evaluation revealed that CelLink is inapplicable only when there are no positive feature correlations between the datasets, a scenario that inherently disrupts biological alignment (Supp. Fig. 9B-C). Beyond its robustness, CelLink is computationally efficient. In the second phase of its pipeline, the number of iterations for unbalanced optimal transport typically ranges from two to ten under the model default settings, resulting in a time complexity of *O*(*N* ^2^) (*N* denotes the batch size). It takes only around 2-5 minutes for CelLink to integrate simulation datasets with 10,000 samples ((Supp. Fig. 9D). Meanwhile, the space complexity of CelLink is also *O*(*N* ^2^).

These characteristics enable CelLink to efficiently integrate large-scale datasets. For instance, it integrated simulated datasets containing 100,000 samples in approximately 25 minutes to 1 hour using an Apple M1 Pro chip (8-core CPU) with 16 GB of RAM. This scalability positions CelLink as a critical tool for generating large-scale paired single-cell multi-omics datasets, a foundational step toward constructing single-cell multi-modal foundation models.

## Discussion

Single-cell multi-omics integration is a pivotal case of machine learning domain adaptation in bioinformatics. Traditionally domain adaptation problems are solved by sample alignment or feature alignment between the source and target data. In the case of single-cell multiomics integration, sample alignment methods typically look for cell-to-cell matching between the two datasets. The cell-to-cell matching is either one-to-one like MaxFuse and Seurat V3 [7, 24], or many-to-many like UniPort and SCOT v2 [25, 6]. However, feature alignment methods transform features from different domains into a common latent space, which demands sufficient samples and related features to effectively capture the underlying patterns of the aligned distributions, like LiGER and scGLUE [4, 24]. These methods face challenges in integrating multi-omics data with weak feature linkage and imbalanced cell populations. Imbalanced cell populations between datasets are common due to the inherent characteristics of datasets and the application of different data quality control standards during the preprocessing of each dataset. These imbalances, along with weak feature linkage, present significant challenges for integrating multi-omics data. Therefore, CelLink is developed based on optimal transport, a classic sample alignment method.

The key issue of imbalanced cell populations and weak feature linkage lies not only in the cell counts but also in the exacerbated imbalance among cell types or cell states, comprising various sub-distributions of the two datasets. Previous optimal transport-based methods do not perform well on this problem because they do not address the varying proportions and complexities of these sub-distributions within the overall distribution [26, 6, 25]. Although Unbalanced Optimal Transport (UOT) incorporates the imbalanced mass between the source and target distributions [12], it still hardly achieves the globally optimal transport map when different sub-distributions in the source and target data partially overlap in cases of weak feature linkage, such as between transcript and protein data.

Motivated by this, we developed CelLink based on two key processes. The first process, similar to MaxFuse, solves the issue of weakly linked features by smoothing the feature profile using neighboring cells and normalizing the feature expression values to enhance the comparability. The second process addresses imbalanced cell populations by performing iterative optimal transport. In this process, cell type or cell state is utilized as the label of sub-distribution to refine the alignment for cells that overlap with other sub-distributions. In this way, CelLink automatically filters out noise cells that cannot be aligned. To reduce the computation cost, CelLink also provides functions to split the data into multiple partitions in low sample sizes.

We compared CelLink with four other baselines using multiple datasets with various omics profiles. We demonstrated that CelLink not only significantly outperformed other methods in scenarios with imbalanced cell populations, but also showed slight improvements in datasets with balanced cell populations. Even without cell-type annotation, the non-iterative pipeline of CelLink still achieved excellent integration performance. Given CelLink’s strong feature imputation capabilities, it largely helps build multi-modality foundation models by generating paired single-cell multi-omics datasets.

## Methods

### The Pipeline of CelLink Model Annotation

Assume the dimension of X as *n*_*x*_ * *n*_*g*_, and the dimension of Y as *n*_*y*_ * *n*_*p*_. Assume the number of linked gene-protein features as *n*_*l*_. Then the linked modality *X*_*l*_ and *Y*_*l*_ are *n*_*x*_ * *n*_*l*_ and *n*_*y*_ * *n*_*l*_. Assume the number of cells that can be potentially aligned as *n*_*c*_. Assume *N* = *max*(*n*_*x*_, *n*_*y*_). The annotations of other variables mentioned in the pipeline are shown in the following table.

**Table 1:** Annotations of the variables and their meanings

| Variable Name | Meaning |
| --- | --- |
| $X, Y$ | The all-feature modalities $X, Y$ |
| $n_x, n_y$ | The number of cells in modality $X, Y$ |
| $n_{mx}, n_{my}$ | The number of matched cells in modality $X, Y$ |
| $n_g, n_p$ | The number of features in modality $X, Y$ |
| $n_l$ | The number of linked features between $X$ and $Y$ |
| $S_x \in (-1, 1)^{n_x * n_x}, S_y \in (-1, 1)^{n_y * n_y}$ | The cell-cell similarity matrices of $X, Y$ |
| $G_x \in \{0, 1\}^{n_x * n_x}, G_y \in \{0, 1\}^{n_y * n_y}$ | The K-nearest-neighbor graphs of $X, Y$ |
| $K_x, K_y$ | The diagonal matrices of the number of neighbor cells for each cell in $X, Y$ |
| $X_l, Y_l$ | The linked-feature modalities of $X, Y$ |
| $\tilde{X}_l, \tilde{Y}_l$ | The smoothed linked-feature modalities $\tilde{X}_l, \tilde{Y}_l$ |
| $D_{xy} \in (0, 2)^{n_x * n_y}$ | The distance matrix between $X_{ss}$ and $Y_{ss}$ |
| $U_x, U_y$ | The final cell-cell correspondence matrix between $X$ and $Y$ |

### Model Input and Preprocessing

The input of CelLink is the all-feature modality and linked-feature modality X, Y. To better differentiate cells of different cell types, we smooth gene expressions and protein marker intensities for each cell based on its neighbor cells which have the highest feature similarity to it. The smoothing process is as follows: (i) Calculate the cell-cell similarity matrices of all-feature X, Y as *S*_*x*_(*n*_*x*_ * *n*_*x*_) and *S*_*y*_(*n*_*y*_ * *n*_*y*_). The default metric to calculate similarity is the Pearson Correlation Coefficient. (ii) Build the K-nearest-neighbor graph for X and Y based on *S*_*x*_ and *S*_*y*_, named *G*_*x*_ and *G*_*y*_, with nodes as cells and edges as the correlation coefficient between cells. (iii) Smooth the linked-feature profile of each cell from its nearest neighbors 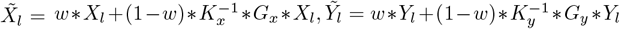, where 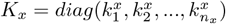, 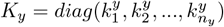, and 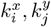 represent the number of neighboring cells for cell *i* and *j*.

#### Data Partition

To better fit the model and save the computational resource, we provide an optional function to split data into multiple partitions so that cell counts between modalities are down-sampled to be closed. For example, we split a CODEX data comprising 120,000 cells into 10 partitions. Each partition will integrate with a scRNA-seq with 10,000 cells.

Each pair of batching of 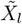 and 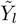 are the inputs of the phase I.

### Phase I: Balanced Optimal Transport

(i) Calculate the cell-pairwise distance matrix *D*_*xy*_(*n*_*x*_ * *n*_*y*_) from *X*_*ss*_ and *Y*_*ss*_. The distance between cell *i* from modality *X* and cell *j* from modality *Y* is 1 − *corr*(*X*_*ss,i*_, *Y*_*ss,j*_).

(ii) Determine *D*_*xy*_ as the cost matrix, infer the original cell-cell correspondence matrix *U*_0_(*n*_*x*_ *n*_*y*_) between *X*_*ss*_ and *Y*_*ss*_ by balanced optimal transport. We assume the mass of each cell as one. Thus, the mass transported from cells in X to Y is the correspondence. The OT problem is expressed as

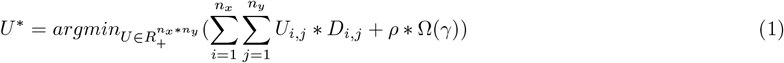

subject to (1) 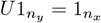 (2) 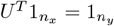 (3) *U*_*i,j*_ ≥ 0 for all *i, j*.

The entropy regularization term 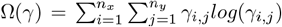 is incorporated by the Sinkhorn algorithm to adjust the sparsity of transport map and accelerate the convergence of the problem [27].

(iii) With the transport map *U*, we predict cell types for each cell by the maximum of weight the cell type from the other modality. The cell-type matched cells are filtered out while the unmatched ones are retained and will be re-aligned in the second phase. The transport map, also known as the original cell-cell correspondence, is stored as *U*_0_.

### Phase II: Iterative Unbalanced Optimal Transport

Iterative unbalanced OT is executed distinctly for unmatched cells within each modality with all cells from the other. Take unmatched cells in modality X as an example. Assume the number of retained unmatched X cells in the *kth* iteration is *n*_*xk*_. The procedure is:

(i) Calculate the cell-pairwise distance matrix between *n*_*xk*_ unmatched cells in X and all cells in Y. The distance metric is the same as phase I.

(ii) Determine the distance matrix as the transport cost. Since our goal is to find the optimal mapping for unmatched cells in X, we allow cells from X to reach their nearest destination in Y by adjusting the mass of Y cells. Thus, we still assume the initial mass of each cell as one, but only enforce that the mass of cells in X remains unchanged while achieving the minimal transport cost through unbalanced optimal transport (UOT). The UOT problem is expressed as

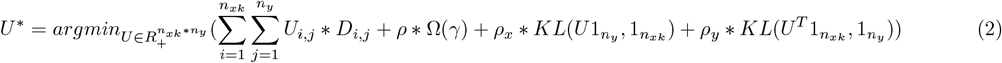

where the entropy regularization term 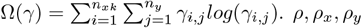 are penalty coefficients of the transport map sparsity, the mass change of X after transport, and the mass change of Y after transport, respectively.

(iii) With the transport map *U*_*k*_ (*k* denotes the current iteration index), we predict cell types for each cell by the maximum of weight the cell type from the other modality. The cell-type matched cells are filtered out while the unmatched ones are retained and will be re-aligned in the second phase. The transport map is stored as *U*_*k*_.

(iv) Repeat (i), (ii), (iii) until the predicted cell types for unmatched cells do not change, indicating the convergence of unbalanced OT and the left cells cannot be aligned.

The optimization of both balanced and unbalanced optimal transport (OT) mentioned above is executed using the Sinkhorn algorithm. This algorithm efficiently produces a relatively dense transport map (cell-cell correspondence matrix). For users requiring sparse results, albeit with much higher computational complexity, we offer an alternative option: skip Phase I and implement iterative UOT with L-BFGS-B optimization [28] in Phase II.

### Phase III: Domain Transfer and Output

We individually impute feature profiles of the other modality for matched cells in X and Y. The procedure to impute Y modality for X is as follows:

(i) Assume the UOT loop runs *t* times. The series of cellular correspondence matrices *U*_0_, *U*_1_, …, *U*_*t*_ is consolidated into a unified matrix *U*_*x*_ based on cell IDs. For every cell in *U*_0_ that does not align correctly, its cell correspondence vector is replaced with the corresponding vector from *U*_*f*_. The subscript *f* stands for the index of the matrix among the series that correctly aligns the previously unmatched cell.

(ii) Assume the number of matched cells in X is *n*_*mx*_, and the dimension of *U* is *n*_*mx*_ * *n*_*y*_. The Y-feature modality for X is imputed as *X*_*imputed*_ = *U* * *Y*.

(iii) To further evaluate the goodness of the integration, we jointly embed *X*_*imputed*_ (*n*_*mx*_ * *n*_*p*_) and *Y* (*n*_*y*_ * *n*_*p*_) into a low-dimensional space and visualize it by UMAP plot.

(iv) The procedure to impute X modality for Y is in the same way, with the final cell-cell correspondence matrix and the X-modality feature for Y denoted as *U*_*y*_(*n*_*my*_ * *n*_*x*_) and *Y*_*imputed*_ (*n*_*my*_ * *n*_*g*_). The output of CelLink is *U*_*x*_, *U*_*y*_ and *X*_*imputed*_, *Y*_*imputed*_.

### Systematic benchmarking of various single-cell multi-omics datasets

We compared CelLink with four other baseline methods, including non-iterative CelLink, MaxFuse, Seurat V3, and SCOT v2. Non-iterative, CelLink, MaxFuse, and SCOT v2 were implemented in Python, and Seurat V3 was implemented in R. We pre-processed all benchmarking datasets in the same way for all methods and used the default hyperparameters suggested by each method. The pre-processing step includes excluding low-quality cells and extracting highly variable genes and proteins for integration. We benchmarked CelLink on seven single-cell multi-omics datasets: three scRNA-seq and CODEX datasets, and four CITE-seq datasets. The evaluation metrics are shown below. We utilized metrics (1), (2), (5), (6) for the datasets with unpaired cells, and (3), (4), (5), (6) for the datasets with paired cells.

### Evaluation Metrics

(1) Average Silhouette Width (ASW): This metric evaluates the global integration performance by measuring the goodness of clustering. For each cell in modality X, it compares its distance to all other cells of the same cell type with the distance to all other cells of the nearest cell type in modality Y in the embedding space. Cells in modality Y are compared with modality X in the same way. ASW is calculated as:

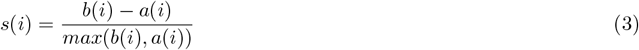

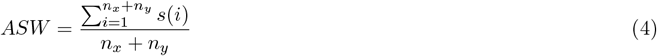

where for cell *i, a*(*i*) is its average distance to all cells in the same cell-type cluster, *b*(*i*) is its smallest average distance to all cells among other cell types and *s*(*i*) is its Silhouette width.

(2) Graph Connectivity Score (GCS): This metric evaluates the local integration performance by measuring the cohesion of data modalities and cell types in the embedding space. It calculates the proportion of the largest connected component within a cell-cell network to the total number of cells. GCS is calculated as:

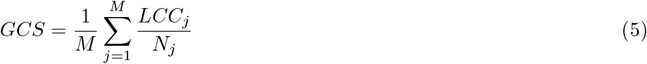

where *M* denotes the number of cell types, *LCC*_*j*_ represents the number of cells in the largest connected component, and *N*_*j*_, is the total number of cells in cell type *j*. The largest connected component is the maximum number of cells that are linked in the *k* − NN graph built from the real and aligned transcriptomic or proteomic expression profile of cells from all modalities.

(3) Feature Imputation accuracy (FIA): This metric evaluates the integration performance by calculating two scores (sub-metrics): (i) the Pearson Correlation coefficient and (ii) the RMSE score between the imputed feature and true feature profiles for each cell in both modalities. In this case, assume *n*_*x*_ = *n*_*y*_ = *n*. They are calculated as:

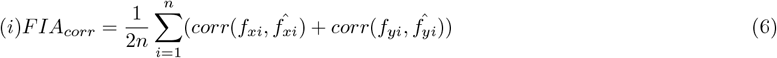

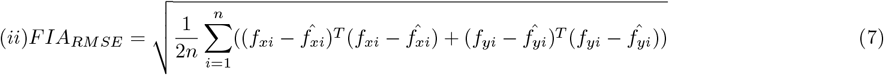

where 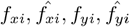 represent the true and imputed feature profile for cell *i* in modality *X* and *Y*, and 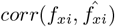 denotes the Pearson Correlation coefficient between them.

(4) Fraction Of Samples Closer Than True Match (FOSCTTM): This metric evaluates the integration performance by determining how many cells from an alternate modality are closer to the target cell than its ground-truth paired cell in the alternate modality. In this case, assume *n*_*x*_ = *n*_*y*_ = *n*. FOSCTTM is calculated as:

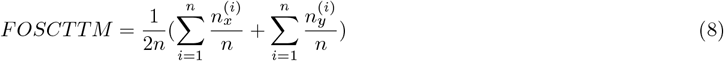

where for each *i*, 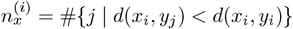 with *d* denoting the Euclidean distance between the embedding of cells.

(5) Cell-type matching accuracy (CTMA): This metric evaluates the integration performance by checking the percentage of cells whose matched cell types are equal to their original cell types. We matched the cell type for each cell in one modality by the cell type that has the maximum correspondence score from the other modality. For a given cell *i*, its correspondence score for cell type *t* is calculated as *Score*_*i,t*_ = Σ_*jint*_ *U* (*i, j*). Then, CTMA is calculated as:

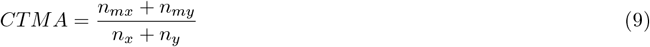

(6) Count of precisely matched cells (CPMC): This metric evaluates the integration performance by calculating the number of cells whose matched cell types are equal to their original cell types. It is a supplementary metric to (5), judging whether a method can succeed in aligning the maximum number of cells. CPMC is calculated as:

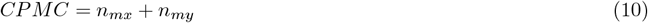

where the meanings of the parameters *n*_*x*_, *n*_*y*_, *n*_*mx*_, *n*_*my*_ are consistent with those provided in the model annotation table. Since cell-type information is utilized as a criterion to determine if each cell is successfully aligned in each iteration, it is natural that CelLink significantly outperforms other methods. Thus, we consider this metric as a supplementary evaluation.

### Cell subtyping

Given the cell subtypes of one data, CelLink performs cell subtype annotations for another data by the inferred cell-cell correspondence matrix. For example, assume scRNA-seq as *X* and CODEX as *Y*. If cell subtypes of *X* are provided, subtypes of cells in *Y* are annotated by the maximum subtype correspondence score in the matrix *U*_*y*_. For a given cell *i*, the subtype correspondence score for subtype *t* is calculated as *Score*_*i,t*_ = Σ_*j*∈*t*_ *U*_*y*_(*i, j*). If cell subtypes of *Y* are provided, subtypes of cells in *X* can be annotated similarly.

### Spatial Cell-cell Communication Analysis

Spatial cellular communication is inferred by assessing spatial proximal ligand-receptor (LR) interactions between specific cell-type pairs relative to random pairs by permutation test using CellChat v2 [20]. This analysis is performed with HPAP024 CODEX data after imputing gene expression profiles from scRNA-seq. The following parameters are set using CellChat: trim=0.1, distance.use = FALSE, interaction.range=250, contact.dependent=TRUE, contact.range=100. Additionally, only communication pathways involving at least 10 cells are retained. LR pairs are considered significant if they show statistical significance by CellChat and have the cell-type level expression of ligand and receptor above 0.01 to avoid noise.

### HPAP scRNA-seq and CODEX datasets analysis

scRNA-seq: The team conducted a comprehensive single-cell RNA sequencing analysis starting with basic quality control a Nextflow-based snRNA-Seq workflow [29] for data processing and downstream analysis using Seurat 4 [13] and other tools described here. STARsolo [30] was used to align the 10X Chromium short read data to the human GRCh38 (primary assembly) genome and mapped to GENCODE v42 annotations. The initial quality filtering was fairly standard, keeping cells with more than 500 features and less than 15% mitochondrial reads while ensuring genes were detected in at least three cells. Identifying real cells versus empty droplets was approached systematically by testing three different methods: EmptyDrops [31] with a strict 1% FDR, a knee-point threshold method, and finally a Dropkick-based threshold approach. The team ultimately selected the Dropkick method [32], applying sample-specific UMI thresholds. Ambient RNA contamination was handled through a two-step SoupX-based process [33]. After testing both automated and manual approaches, we settled on a hybrid method: first using automatic contamination estimation, followed by a manual adjustment with an additional 0.2 contamination fraction. Doublet detection relied on scDoubletFinder [34] with a sample-based increase of the doublet rate parameter up to a value of 0.3, which on average identified 16% of cells as potential doublets, which were then removed. A manual inspection was performed to remove cell clusters single-donor derived. Integration relied on Harmony [35] and included the following covariates: HPAP ID, technical platforms, and tissue source. For clustering, we used 2,000 most variable genes, using 15 principal components and a Louvain algorithm with 0.3 resolution. The final dataset includes key clinical metadata such as age, disease type, and T1D-related groupings. CODEX: HPAP OCT tissue blocks were sectioned at 8 μm and prepared for CODEX imaging according to the PhenoCycler User Manual (Akoya Biosciences). PhenoImager (Akoya Biosciences) acquired image tiles and performed tile stitching, deconvolution, background subtraction, and image registration.

Processed images were then segmented into cell objects using HALO AI (Indica Labs) trained on manually annotated DAPI images. Cells were exported as tables with associated cell marker intensity data. These cell tables were further processed by Pearson residual transformation followed by Principal Component Analysis (PCA). The resulting principal components were used as input for Uniform Manifold Approximation and Projection (UMAP), reducing the number of available dimensions to two. These two dimensions were used in Leiden clustering, which resulted in a list of clusters and Z-score normalized average intensities for each cell marker. These clusters and cell marker intensities were visualized as a heatmap, which was then inspected for cell marker intensity signatures that could identify the type of cell corresponding to that cluster. Clusters that contained conflicting cell marker data were sub-clustered to identify less common cell populations. Cells were then assigned cell types according to their cluster membership.

### Pre-process of the scRNA-seq, CODEX, and CITE-seq datasets

Among the three scRNA-seq & CODEX datasets (HPAP023, HPAP071, Tonsil) and four CITE-seq datasets (two balanced and two imbalanced), we applied the following preprocessing steps. For the transcriptomic data, gene expressions were normalized using the log transformation, and the top 5,000 highly variable genes were selected. Subsequently, we excluded non-static genes with standard deviations exceeding 0.05. For the CODEX data, protein marker intensities were normalized using the log transformation, and filtered to remove non-static markers with standard deviations above 0.05. Finally, the linked-feature profiles from the two modalities were further scaled for integration. For the additional scRNA-seq & CODEX datasets (HPAP024 and Bone Marrow), the same preprocessing steps were applied, except that the top 10,000 highly variable genes were selected instead.

### Pseudocode for the pipeline for generating simulation datasets

The pseudocode to generate simulation datasets is shown below. For step 11 in the pseudocode, the mathematical rationale to adjust **x**_d2 linked_ so that for each of the linked feature *i* = 1, …, *n*_linked_,

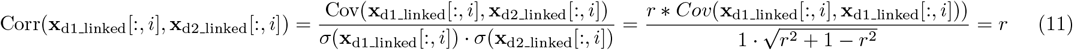

#### Algorithm 1

Simulation Pipeline for Generating Multimodal Data

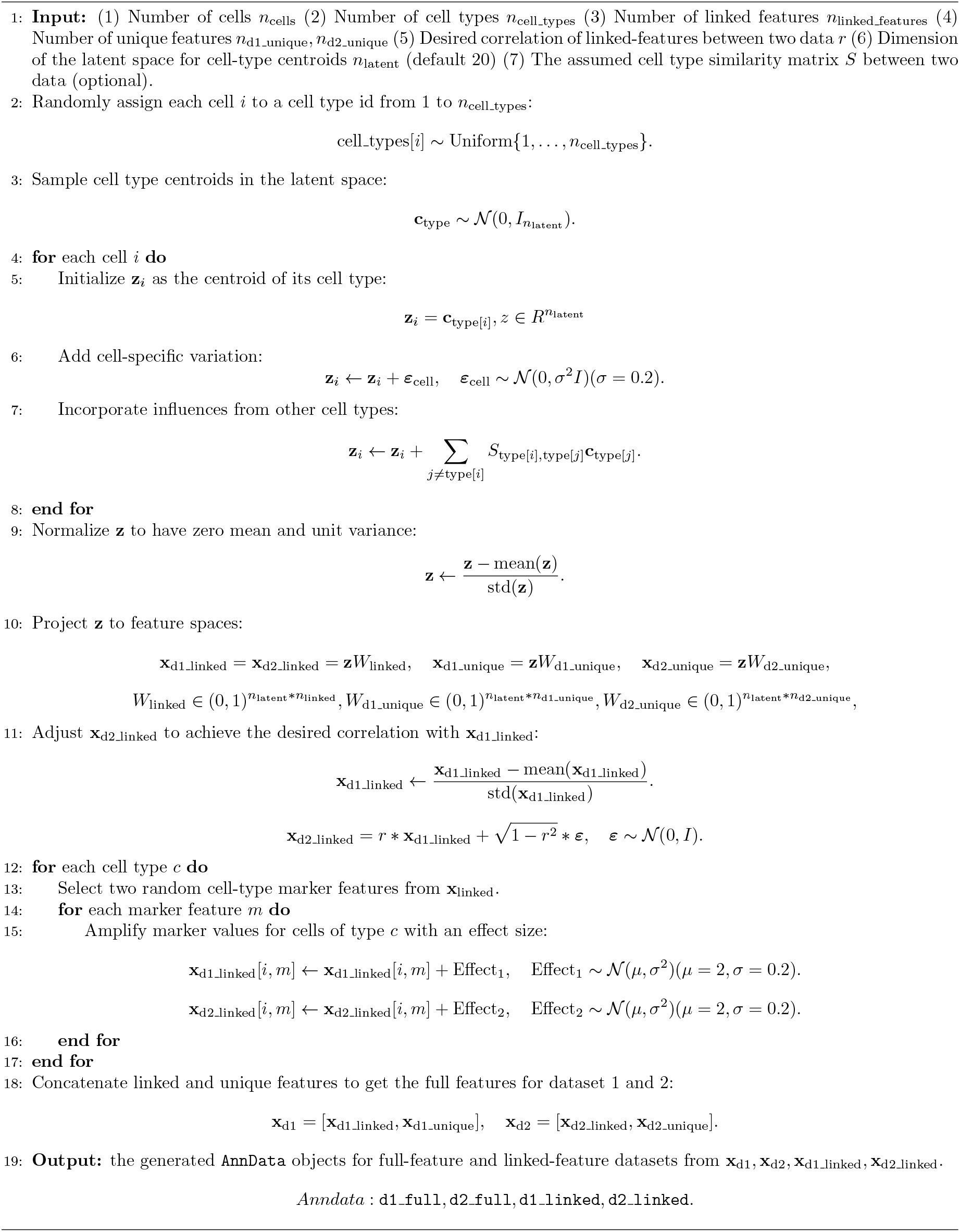

## Data availability

All datasets used in this study are publicly available. The three scRNA-seq and CODEX datasets of HPAP donors come from PANC-DB [2] with link https://hpap.pmacs.upenn.edu/explore/download?matrix. The CITE-seq PBMC data is from Hao et al.[13]: https://atlas.fredhutch.org/data/nygc/multimodal/pbmc_multimodal.h5seurat. The CITE-seq BMC data is from Hao et al.[13]: https://satijalab.org/seurat/articles/multimodal_reference_mapping.html (file: ‘bmcite’ with ‘SeuratData’). The CODEX fata of tonsil tissue is from Kennedy et al.[11]: https://onlinelibrary.wiley.com/doi/10.1002/eji.202048891. The scRNA-seq data of tonsil tissue is from King et al.[36]: https://www.ncbi.nlm.nih.gov/geo/query/acc.cgi?acc=GSE165860. The bone marrow scRNA-seq and CODEX datasets are from Bandyopadhyay et al. [14], with scRNA-seq: https://www.ncbi.nlm.nih.gov/geo/query/acc.cgi?acc=GSE253355, and CODEX: https://doi.org/10.25452/figshare.plus.c.7174914.

## Code availability

All code used in this study, including the CelLink software and experiments, can be found at our GitHub repository: https://github.com/liu-bioinfo-lab/CelLink. Additionally, CelLink is published as a Python package on PyPI for easy download and installation.

