## Supplementary Materials for "CelLink: integrating single-cell multi-omics data with weak feature linkage and imbalanced cell populations"

### Supplementary Notes

#### Details about the intuitions to design iterative optimal transport with Sinkhorn optimization

**Intuition to utilize Sinkhorn algorithm as the optimization method for optimal transport.** The optimal transport algorithm has various optimization methods, including Linear Programming, Sinkhorn Algorithm, L-BFGS, .etc [1, 2, 3]. Among these, the Sinkhorn algorithm emerges as the predominant choice for OT in single-cell multi-omics integration for its efficiency. This preference is underpinned by two principal factors. To illustrate the implementation of Sinkhorn, its pseudocode is provided below.

First, the Sinkhorn algorithm incorporates an entropy term into the loss function, which regulates the sparsity of the cell-cell correspondence matrix. The introduction of this entropy loss is pivotal as it renders the loss function strictly convex, thereby facilitating the convergence to a unique solution. This characteristic is crucial in ensuring that the iterative alignment process is both stable and reliable.

Second, the Sinkhorn algorithm uniquely addresses the scalability challenges inherent in integrating large-scale datasets, which is a common scenario in single-cell studies. Unlike other optimization methods, which typically demand substantial memory and computational resources, the Sinkhorn method maintains efficiency even as the number of cells escalates.

**Intuition to utilize the iterative alignment strategy.** Although Sinkhorn is an efficient method, its entropy term, while beneficial for achieving rapid convergence, may lead to the algorithm settling at a local optimum rather than the global optimum. Consequently, this could result in suboptimal alignment of some cells. Such partial overlap, particularly when optimization is trapped in a local optimum, can exacerbate misalignment issues and compromise the accuracy of cell-type alignment. To address this challenge, we developed an iterative optimal transport (OT) strategy that dynamically adjusts the marginal distributions $(a, b)$ and the cost matrix $D$ of the input datasets by modifying the sample populations iteratively. Initially, each cell was assigned a mass of 1 unit in both datasets for fairness. Unbalanced OT was then applied in each iteration to enable flexible alignment, allowing for adjustments in cell mass post-alignment without imposing strict constraints. This approach iteratively removes the matched cells and re-aligns the unmatched cells until the cell-cell matching results stabilize.

Optimal transport with Sinkhorn algorithm:

$$U^* = argmin_{U \in R_+^{n_x * n_y}} (\sum_{i=1}^{n_x} \sum_{j=1}^{n_y} U_{i,j} * D_{i,j} + \rho * \Omega(\gamma)) \tag{1}$$

subject to (1) $U 1_{n_y} = a$ (2) $U^T 1_{n_x} = b$ (3) $U_{i,j} \geq 0$ for all $i, j$.
In the equation, $D$ represents the cost (distance) matrix between cells in the two datasets. The entropy regularization term $\Omega(\gamma) = \sum_{i=1}^{n_x} \sum_{j=1}^{n_y} \gamma_{i,j} log(\gamma_{i,j})$ is incorporated to adjust the sparsity of transport map and accelerate the convergence of the problem [2]. $\rho$ is the penalty coefficient of the transport map sparsity.

**Algorithm 1** Sinkhorn-Knopp Algorithm for Entropic Regularized Optimal Transport

**procedure** $U^* =$ SinkhornKnopp($D, a, b, \rho$, numItermax, tol)
  $K \leftarrow \exp(-D/\rho)$ ▷ Kernel matrix
  $u \leftarrow \mathbf{1}_n$ ▷ Initial scaling vector for rows
  $v \leftarrow \mathbf{1}_m$ ▷ Initial scaling vector for columns
  **for** iter = 1 to numItermax **do**
    $u \leftarrow a/(Kv)$ ▷ Update u
    $v \leftarrow b/(K^\top u)$ ▷ Update v
    **if** $\max(\|a - Kv\|_1, \|b - K^\top u\|_1) <$ tol **then**
      **break** ▷ Convergence check
    **end if**
  **end for**
  **return** $\text{diag}(u) K \text{diag}(v)$ ▷ Compute the transport matrix
**end procedure**

Hint for CellLink pipeline usage: (1) If cell annotation or cell state is provided, BOT combining iterative UOT is suggested. (2) If cell annotation or cell state is not provided, a one-time reciprocal UOT is suggested. (3) If a sparse alignment result is needed, CellLink offers an L-BFGS optimization option [3]. However, this method is significantly more time-consuming, and we recommend limiting the batch size to no more than 5000 when utilizing this approach.

### Supplementary Figures

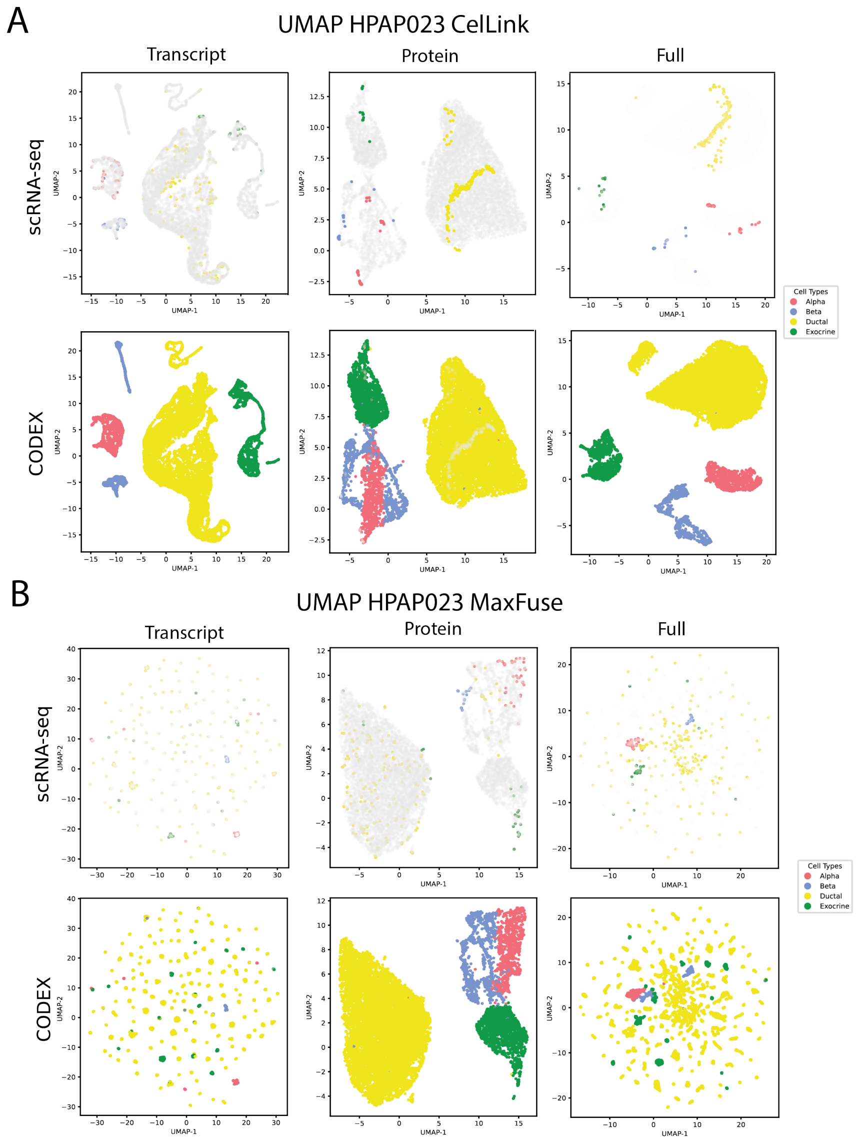


**Supplementary Figure 1. Benchmarking of CellLink on scRNA-seq and CODEX datasets of an AAB donor HPAP023 with MaxFuse from UMAP plots. A.** The UMAP plot exhibits the intuitive integration results from CellLink on HPAP023 donors with the transcript, protein, and full (transcript & protein) profiles. For the transcript profile, scRNA-seq data refers to the original profile while CODEX refers to the the imputed one. For the protein profile, scRNA-seq data refers to the imputed profile while CODEX refers to the original one. **B.** The UMAP plot exhibits the intuitive integration results from MaxFuse on HPAP023 donors with the transcript, protein, and full (transcript & protein) profiles.

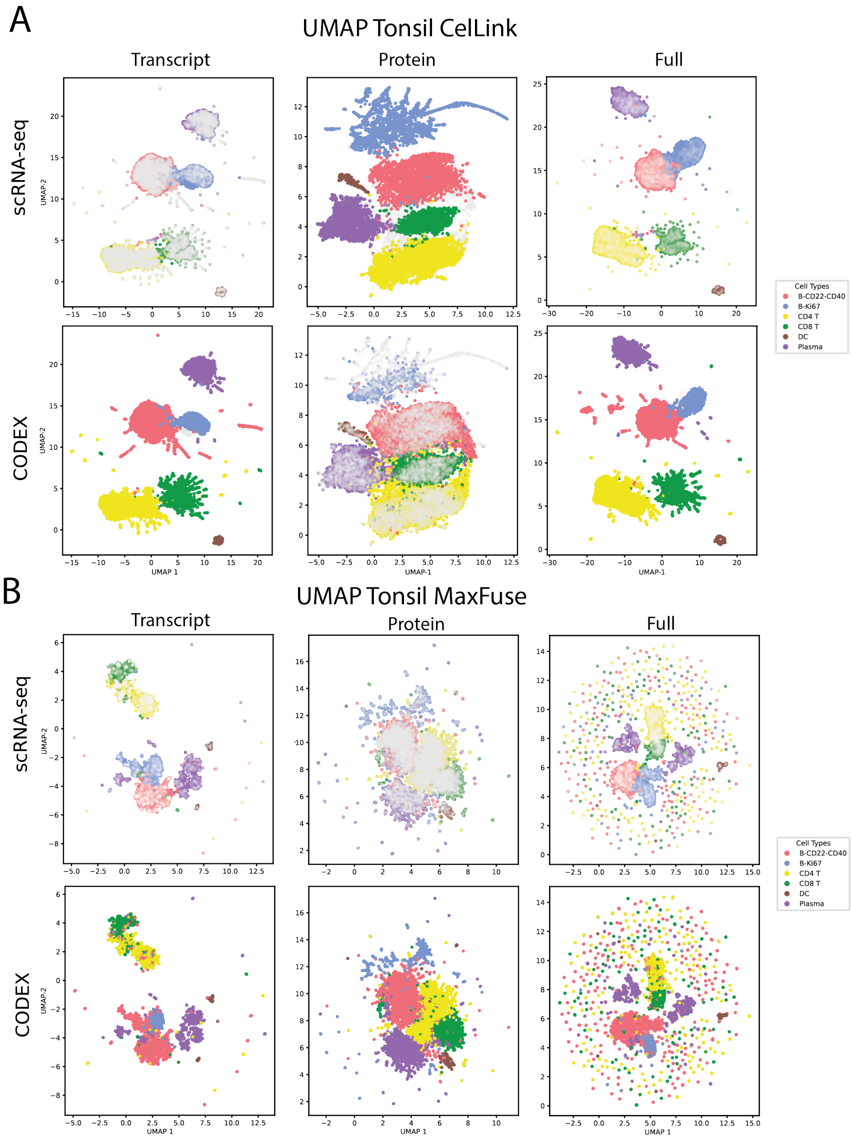


**Supplementary Figure 2. Benchmarking of CellLink on scRNA-seq and CODEX datasets from a tonsil tissue with MaxFuse from UMAP plots. A.** The UMAP plot exhibits the intuitive integration results from CellLink on a tonsil tissue with the transcript, protein, and full (transcript & protein) profiles. For the transcript profile, scRNA-seq data refers to the original profile while CODEX refers to the the imputed one. For the protein profile, scRNA-seq data refers to the imputed profile while CODEX refers to the original one. **B.** The UMAP plot exhibits the intuitive integration results from MaxFuse on a tonsil tissue with the transcript, protein, and full (transcript & protein) profiles.

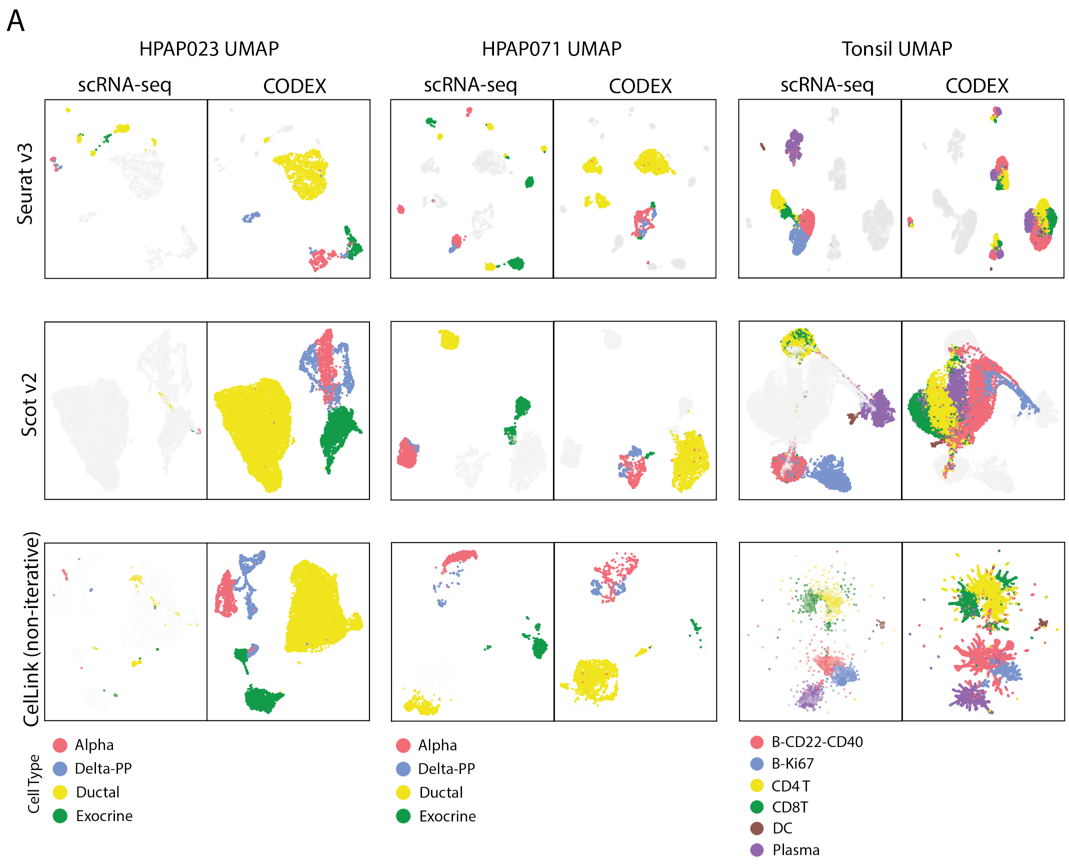


**Supplementary Figure 3. Benchmarking of CelLink on scRNA-seq and CODEX datasets with three baseline methods from UMAP plots. A.** The UMAP plot exhibits the intuitive integration results of three other baseline methods from three datasets with the transcript, protein, and full (transcript & protein) profiles. For the transcript profile, scRNA-seq data refers to the original profile while CODEX refers to the the imputed one. For the protein profile, scRNA-seq data refers to the imputed profile while CODEX refers to the original one.

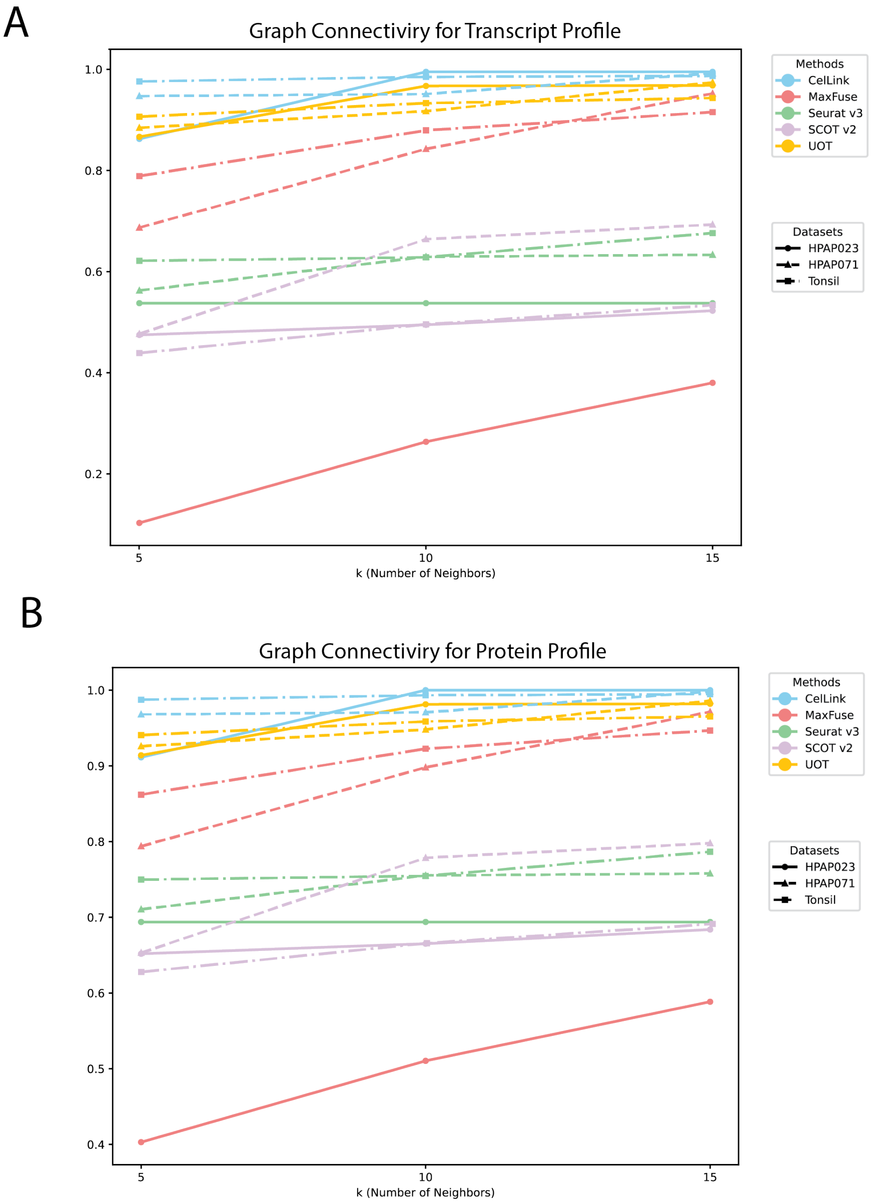


**Supplementary Figure 4.** Benchmarking of CelLink on scRNA-seq and CODEX datasets from GCS. **A.** The line plot displays the graph connectivity scores for CelLink and four other baselines across three scRNA-seq and CODEX datasets, using the transcript profile. Each line indicates the trend of the score when the edges of the graph are built with the increasing number of neighboring cells. The color of the line represents the method. The line type and point shape represent the dataset. **B.** The line plot displays the graph connectivity scores for CelLink and four other baselines across three scRNA-seq and CODEX datasets, using the protein profile.

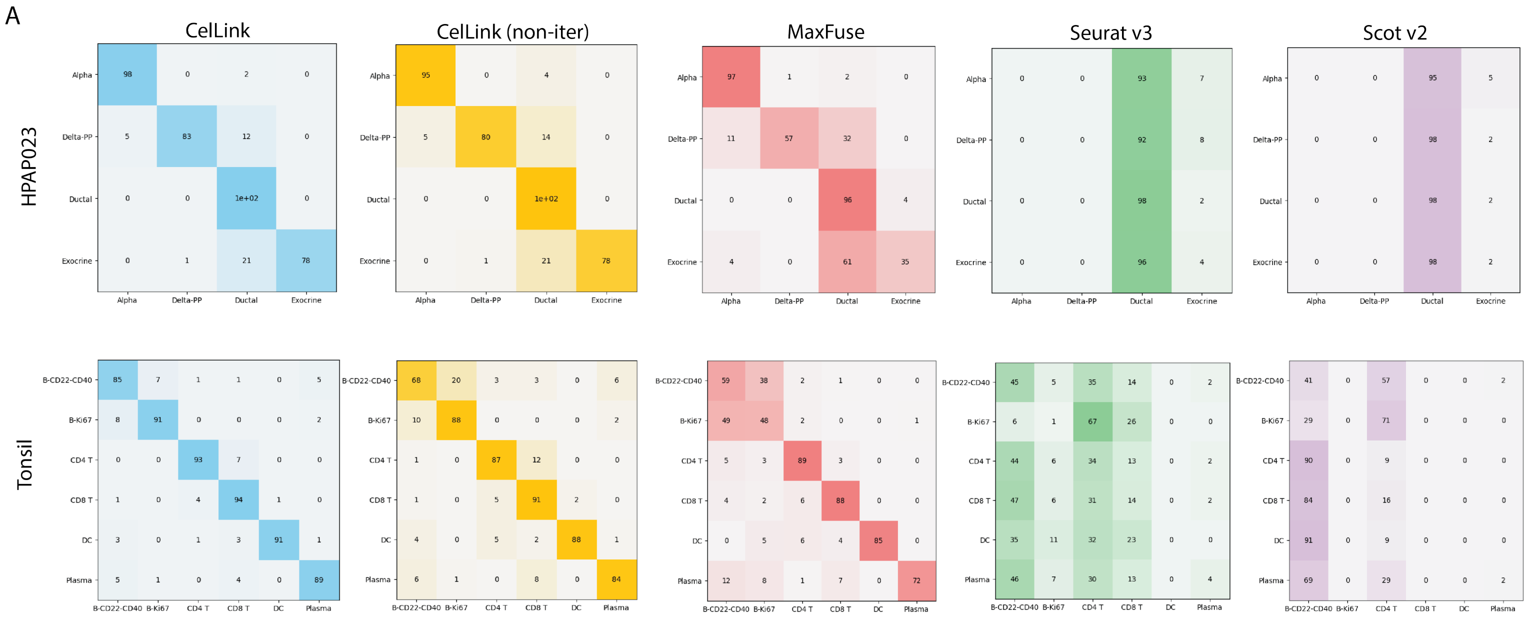


**Supplementary Figure 5. Benchmarking of CelLink on scRNA-seq and CODEX datasets from cell-type matching accuracy. A.** The heatmaps display the detailed cell-type matching result for CelLink and four other baselines on the HPAP023 and Tonsil datasets.

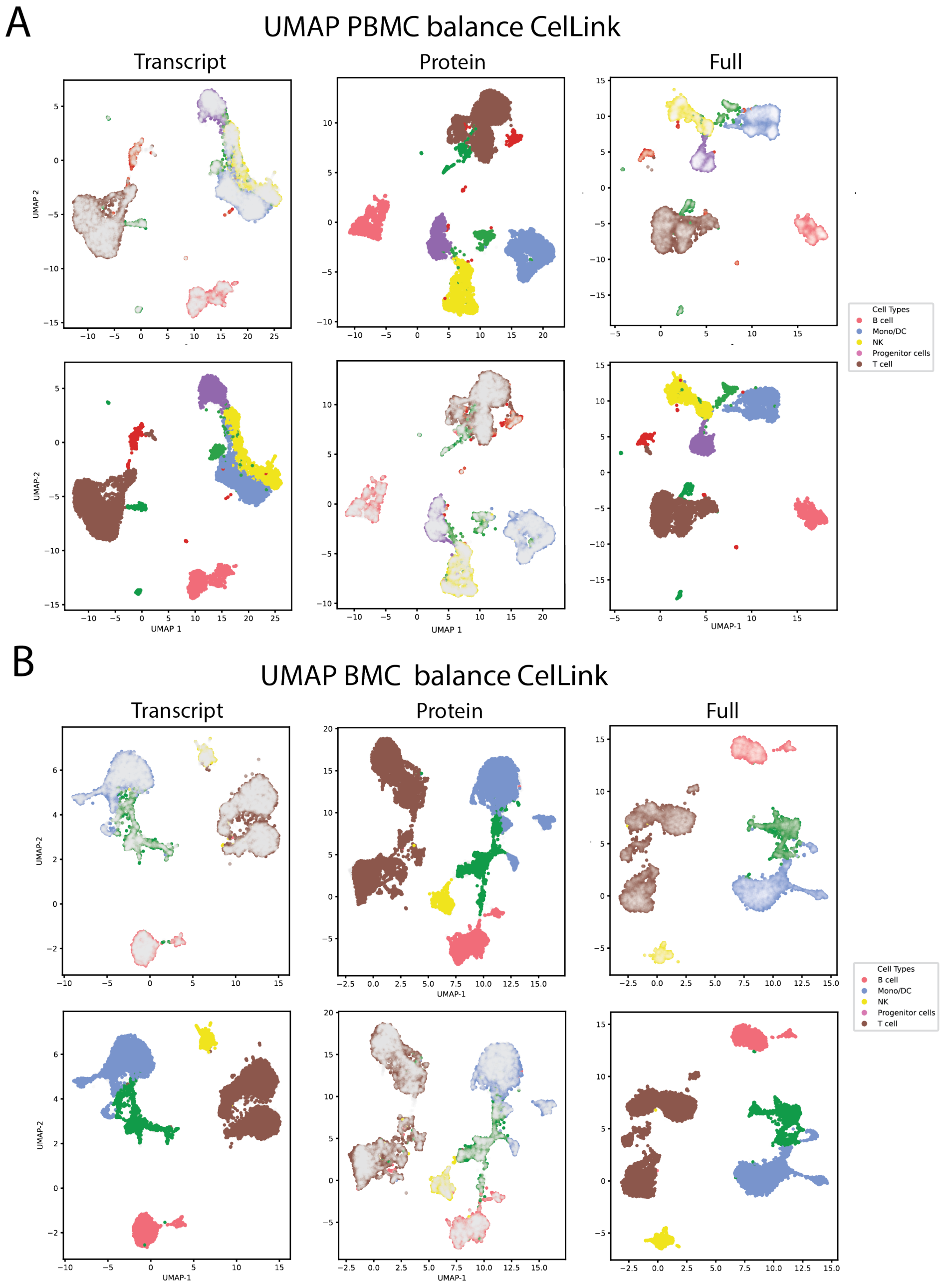


**Supplementary Figure 6. Benchmarking of CelLink on CITE-seq datasets from UMAP plots. A.** The UMAP plot exhibits the intuitive integration results from CelLink on PBMC (balance) data with the transcript, protein, and full (transcript & protein) profiles. **B.** The UMAP plot exhibits the intuitive integration results from CelLink on BMC (balance) data with the transcript, protein, and full (transcript & protein) profiles.

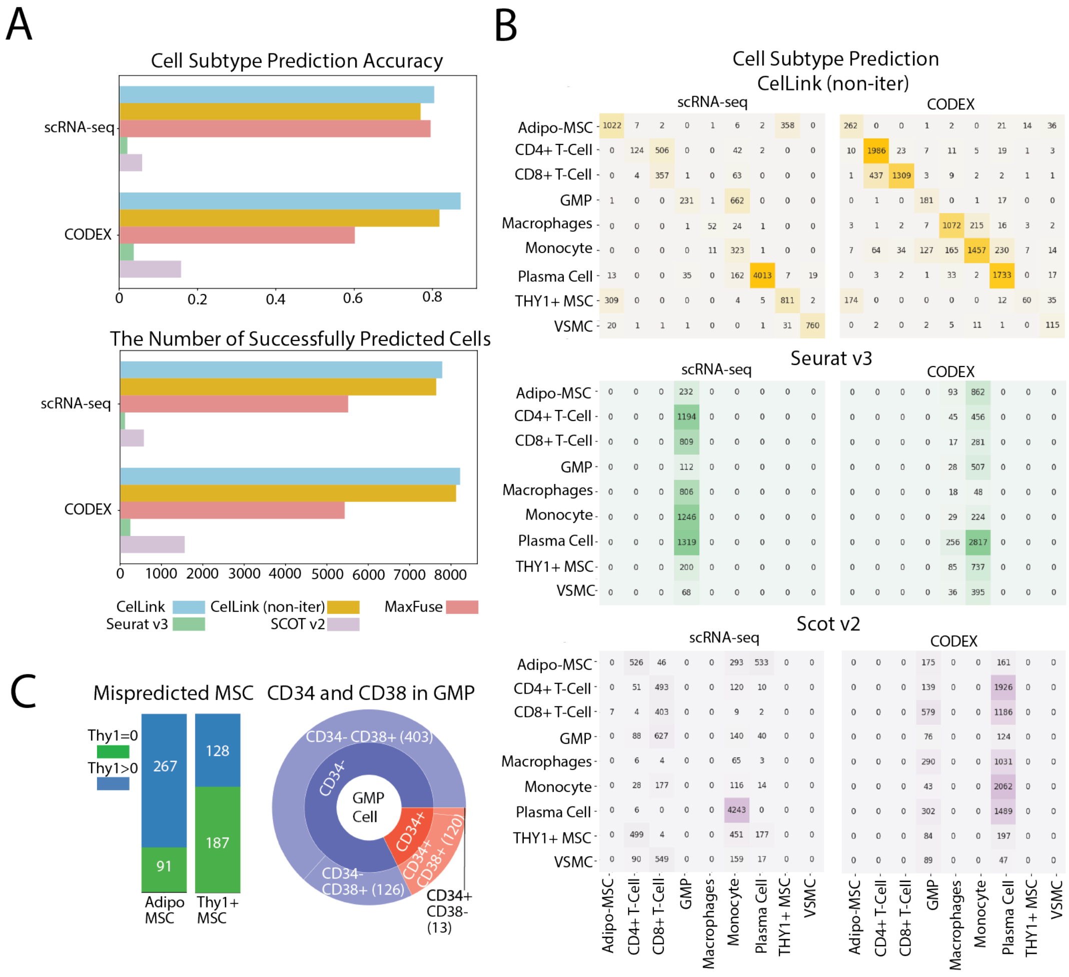


**Supplementary Figure 7. CelLink enables cell subtyping and identifies mislabeled cells. A.** Two horizontal barplots display the cell-type matching accuracy and the number of precisely matched cells for the five methods on the bone marrow dataset. **B.** The heatmaps exhibit the prediction performance of cell subtypes by three baseline methods CelLink (non-iterative), Seurat v3, and SCOT v2. **C.** Left: A barplot depicts the number of cells that express or do not express the Thy1 gene in Adipo MSC and Thy1+ MSC cells. Right: The sunburst chart displays the distribution of CD34 and CD38 coding genes among the mispredicted GMP Cells. Each partition described the number of cells with a distinct combination of CD34 and CD38 marker expressions.

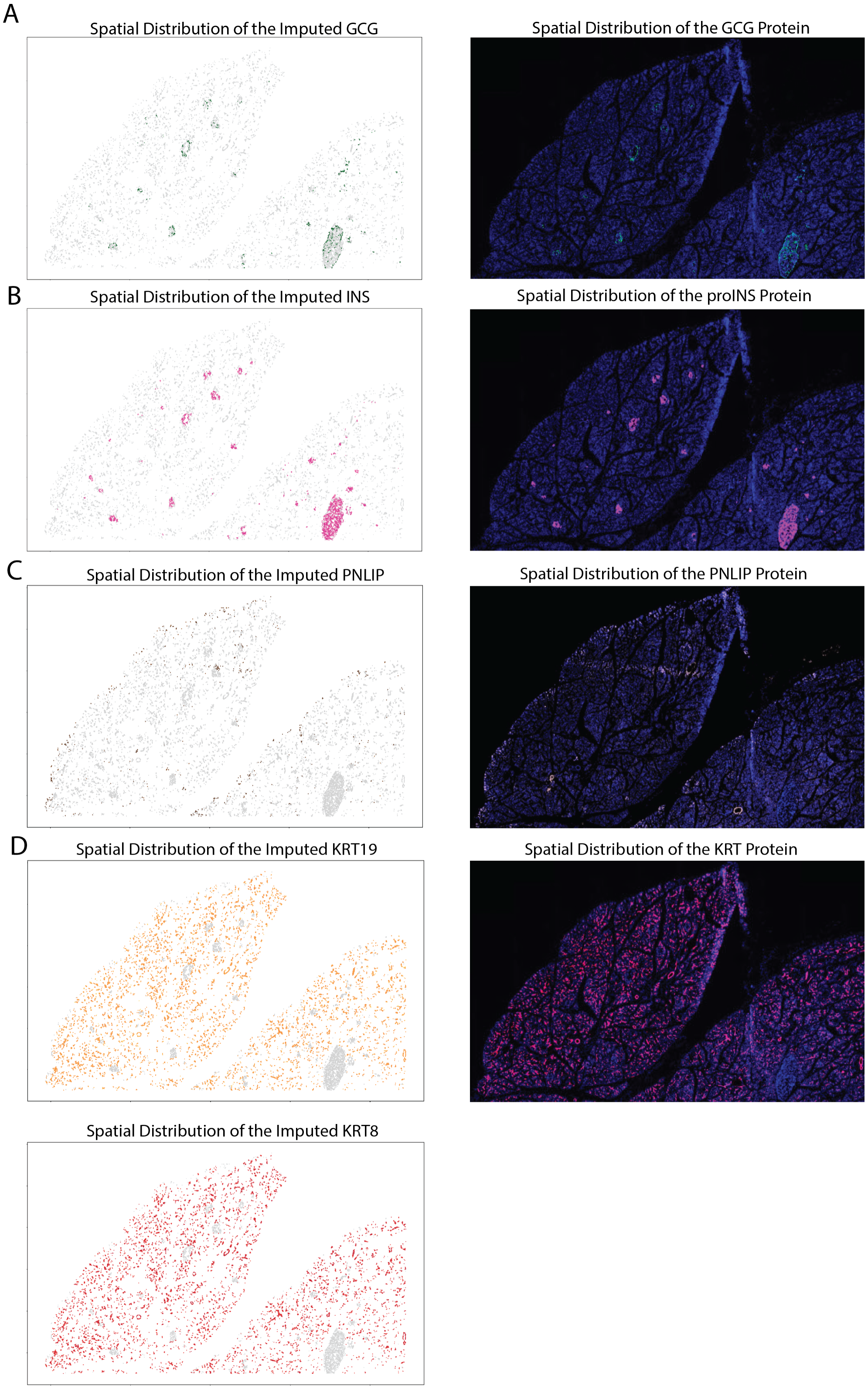


**Supplementary Figure 8. Spatial distributions of the imputed marker gene expressions, original marker protein intensities, and cell neighborhoods.** Protein intensity figures on the right are visualized using Pancreatlas, a specialized website that aggregates and shares data from various human pancreas samples. Some spatial regions in the left-side figures are missing due to the difficulties of the cell segmentation process. A. Left: Spatial distribution of the imputed GCG gene expressions. Right: Spatial distribution of the protein GCG marker intensities. B. Left: Spatial distribution of the imputed INS gene expressions. Right: Spatial distribution of the protein proINS marker intensities. C. Left: Spatial distribution of the imputed PNILP gene expressions. Right: Spatial distribution of the protein PNILP marker intensities. D. Left top: Spatial distribution of the imputed KRT7 gene expressions. Left bottom: Spatial distribution of the imputed KRT8 gene expressions. Right top: Spatial distribution of the protein KRT marker intensities.

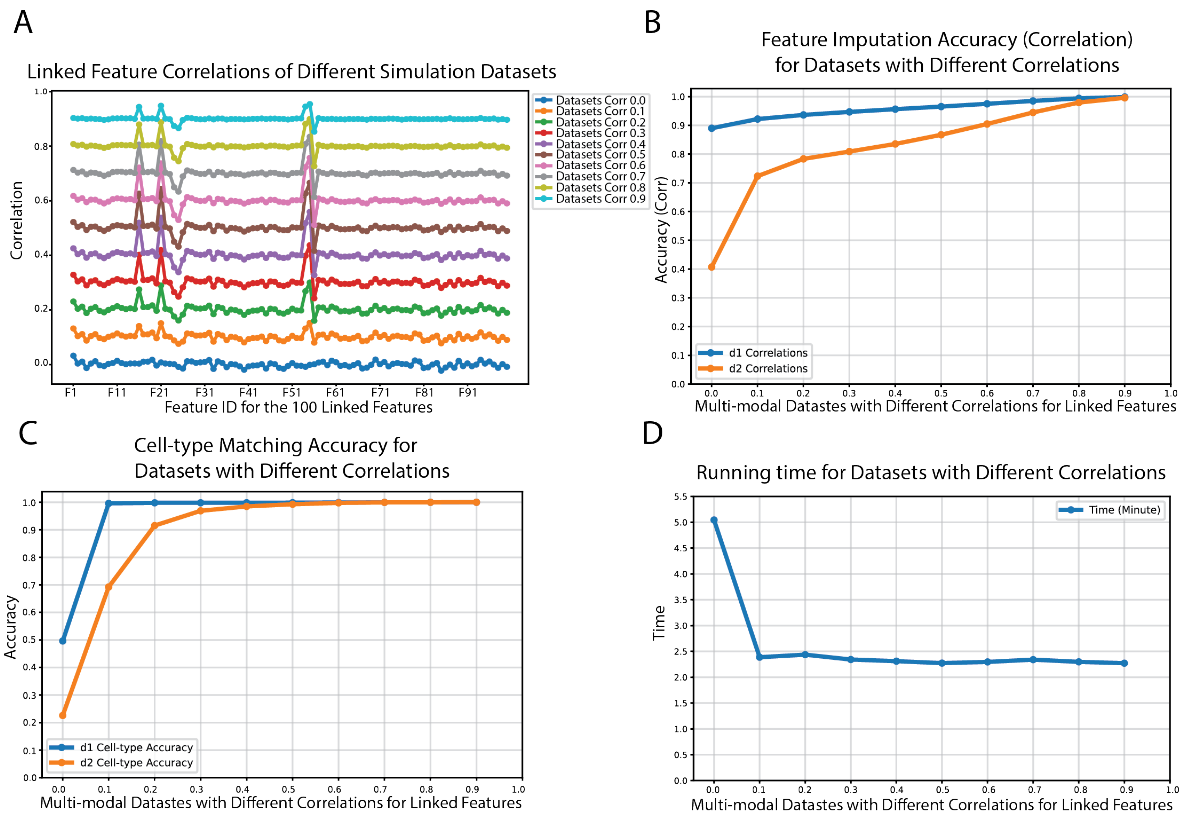


**Supplementary Figure 9. Evaluation of CelLink on large-scale simulation datasets**. **A.** The line plot depicts the correlation between 100 generated linked features with IDs through F1 to F100 between d1 and d2 for integration. Ten simulation datasets, each with d1 and d2, are generated with linked-feature correlations ranging from 0 to 0.9 in increments of 0.1. **B.** The feature imputation accuracy (correlation score in evaluation metric (3)) for d1 and d2 among 10 simulation datasets with different levels of linked-feature correlations. **Hint:** Since CelLink automatically filters unmatched cells, only matched cells are utilized to calculate the feature imputation score. **C.** The cell-type matching accuracy for d1 and d2 among 10 simulation datasets with different levels of linked-feature correlations. **D.** The running time of CelLink on 10 simulation datasets with different levels of linked-feature correlations.

### Supplementary Tables

| Cell Type | scRNA-seq Composition | CODEX Composition |
|---|---|---|
| Ductal | 196 | 7655 |
| Alpha | 36 | 730 |
| Exocrine | 26 | 1455 |
| Delta-PP | 20 | 802 |
| Stellate | 19 | 0 |
| Endothelial | 16 | 3549 |
| Beta | 1 | 0 |
| Immune | 0 | 1823 |
| Pericyte or Smooth Muscle Cell | 0 | 1969 |
| Smooth Muscle Cell | 0 | 372 |
| Neuronal Cell | 0 | 15 |

**Table 1:** Cell compositions across scRNA-seq and CODEX datasets in HPAP023. Only cell types with counts greater than 20 in both datasets are considered. Alpha, Ductal, Delta-PP and Exocrine cells are selected for integration.

| Cell Type | scRNA-seq Composition | CODEX Composition |
|---|---|---|
| Alpha | 629 | 233 |
| Exocrine | 578 | 40 |
| Ductal | 432 | 1097 |
| Delta-PP | 104 | 146 |

**Table 2:** Cell compositions across scRNA-seq and CODEX datasets in HPAP071. Only Alpha, Exocrine, Ductal, and Delta-PP cells are extracted from both datasets. For this analysis, a single partition of the CODEX data (batch size of 1,500) was extracted to evaluate Cellink on a smaller sampled dataset.

| Cell Type | scRNA-seq Composition | CODEX Composition |
|---|---|---|
| B-CD22-CD40 | 2809 | 3262 |
| CD4 T | 2420 | 2997 |
| B-Ki67 | 2245 | 481 |
| Plasma | 1357 | 1590 |
| CD8 T | 1045 | 1540 |
| DC | 124 | 130 |

**Table 3:** Cell compositions across scRNA-seq and CODEX datasets in Tonsil.

| Cell Type | scRNA-seq Composition | CODEX Composition |
|---|---|---|
| CD4 T | 2300 | 263 |
| Mono | 400 | 2633 |
| B | 100 | 742 |
| CD8 T | 100 | 1432 |
| NK | 100 | 1046 |
| other T | 100 | 314 |
| DC | 50 | 184 |
| other | 50 | 186 |

**Table 4:** Cell compositions across split CITE-seq dataset in PBMC Imbalanced.

| Cell Type | scRNA-seq Composition | CODEX Composition |
|---|---|---|
| T cell | 4200 | 498 |
| Mono/DC | 2400 | 254 |
| Progenitor cells | 900 | 164 |
| NK | 330 | 84 |
| B cell | 170 | 1000 |

**Table 5:** Cell compositions across split CITE-seq dataset in BMC Imbalanced.

| Cell Type | scRNA-seq Composition | CODEX Composition |
|---|---|---|
| Lymphoid | 5348 | 5621 |
| Mesenchymal | 2529 | 617 |
| Myeloid | 1308 | 3626 |
| Muscle | 815 | 136 |

**Table 6:** Cell state (lineage) compositions across scRNA-seq and CODEX datasets in Bone Marrow.

| Cell Type | scRNA-seq Composition | CODEX Composition |
|---|---|---|
| Plasma Cell | 4249 | 1791 |
| Adipo-MSC | 1398 | 336 |
| THY1+ MSC | 1131 | 281 |
| GMP | 895 | 200 |
| VSMC | 815 | 136 |
| CD4+ T-Cell | 674 | 2065 |
| CD8+ T-Cell | 425 | 1765 |
| Monocyte | 335 | 2105 |
| Macrophages | 78 | 1321 |

**Table 7:** Cell subtype compositions across scRNA-seq and CODEX datasets in Bone Marrow.

| Cell Type | scRNA-seq Composition | CODEX Composition |
|---|---|---|
| Exocrine | 877 | 535 |
| Ductal | 217 | 8875 |
| Alpha | 55 | 683 |
| Beta | 54 | 2258 |
| Stellate | 38 | 0 |
| Endothelial | 13 | 3285 |
| Delta-PP | 12 | 96 |
| Immune | 0 | 2931 |
| Pericyte or Smooth Muscle Cell | 0 | 2085 |
| Mast Cell | 0 | 430 |

**Table 8:** Cell compositions across scRNA-seq and CODEX datasets in HPAP024. Only cell types with counts greater than 20 in both datasets are extracted. Alpha, Beta, Ductal, and Exocrine cells are selected for integration.
